# Dynamic tracking of social variables in simultaneous brain recordings from socially interacting monkeys

**DOI:** 10.64898/2026.03.12.711496

**Authors:** Sini Simon, Debdyuti Bhadra, Shubhankar Saha, Surbhi Munda, Jhilik Das, SP Arun

## Abstract

Primates are deeply social, but understanding the neural basis has been challenging since it has previously only been possible to record from single monkeys interacting with a social partner. Here, we performed simultaneous wireless neural recordings from two macaque monkeys interacting socially in a natural, unconstrained setting. Neural activity in each monkey encoded key social variables such as allogrooming state, partner identity and joint movements. Interestingly, neurons in the high-level visual cortex of each monkey continuously tracked a social favor signal, i.e. net duration of allogrooming given versus received, providing a neural basis for grooming reciprocity. In each monkey, neural activity was predicted better by his partner’s joints and neural activity, particularly while giving compared to receiving grooming. Thus, the receiver of grooming drove the social interaction, not the groomer. Taken together, our findings elucidate the rich and dynamic neural basis of primate social interactions in an ecologically valid setting.

**SIGNIFICANCE STATEMENT:** Primates are deeply social, but understanding this requires recording neural activity simultaneously and unobtrusively from two animals without constraints. Here, we performed the first-ever simultaneous wireless brain recordings from two monkeys socially interacting in a natural setting. We found that both monkeys’ brains tracked important social variables, such as a social favor signal as well as their partner’s identity and movements. Interestingly, the receiver of grooming seemed to drive the social interaction, as evidenced by referential gestures during behavior, stronger tracking of the grooming receivers’ joints and stronger prediction of the groomer’s brain activity by the receiver’s brain activity. Our results represent an important first step towards understanding primate social interactions in an ecologically valid setting.

## INTRODUCTION

Primates are deeply social and unsurprisingly, social interactions evoke widespread brain activity (Sliwa and Freiwald, 2017; Shepherd and Freiwald, 2018; Wittmann et al., 2018; Gilbert et al., 2021; Deen et al., 2023; McMahon and Isik, 2023). However, understanding the neural basis of social interactions has been technically challenging, because brain recordings are often feasible only under highly artificial, restrained conditions (Fan et al., 2021). Under these conditions, brain activity has been recorded from monkeys interacting with social partners, showing tracking of the partners’ actions, reward states and strategies (Yoshida et al., 2012; Ong et al., 2021; Haroush and Williams, 2015; Chang et al., 2013; Báez-Mendoza et al., 2021). Likewise, brain activity has been recorded simultaneously from socially interacting monkeys, showing interbrain synchrony (Tseng et al., 2018; Gilbert et al., 2021) and also modulation by the partner’s actions (Fujii et al., 2007, 2009). Similar observations have been made in mice (Kingsbury et al., 2019; Zhang et al., 2025), bats (Zhang and Yartsev, 2019; Rose et al., 2021) and even humans (Stephens et al., 2010; Kinreich et al., 2017). With advances in wireless neural recording technology, brain activity has been recorded under increasingly unrestrained and more natural conditions (Roy and Wang, 2012; Schwarz et al., 2014; Berger et al., 2020; Hansmeyer et al., 2023). These advances have prompted calls for concerted efforts to study brain activity in natural and social behaviors (Testard et al., 2021; Cisek and Green, 2024; Parodi et al., 2025).

Despite these advances, several fundamental questions remain regarding our understanding of primate social interactions. First, since primate social interactions are often dyadic with extended bouts of allogrooming, how does each individual keep track of these interactions and know when to switch from receiving to giving grooming? One possibility is that each monkey maintains a social ledger that tracks the net amount of grooming given versus received. Indeed, in a recent study using brain recordings from an individual monkey interacting with a social partner, population neural activity in inferior temporal and prefrontal regions was able to predict several grooming reciprocity signals, suggesting that brain activity can track such long-term social variables (Testard et al., 2024). However, these observations are not fully interpretable without simultaneous brain recordings from both interacting monkeys. In particular, a simple change in the sensory or motor or task-related activity in one individual could easily be mistaken for a social ledger signal since grooming always involves changes in many other variables. However, since one individual giving a social favor always entails the other individual receiving it, a social ledger signal must necessarily be anti-correlated in the brains of the two interacting individuals. Furthermore, the social ledger signal could be tracked as either a social favor or a social debt: the former would increase when giving grooming and decrease when receiving grooming, whereas the latter would show the opposite trend. Resolving this issue will require simultaneous brain recording from both socially interacting individuals.

Second, social interactions involve close coordination between the two individuals, which one of them drives the interaction, deciding where and when to give or receive grooming? The interaction could be driven always by the socially dominant individual, or the interaction could be led each time, by the giver of grooming, or by the receiver of grooming. In the wild, the receiver of grooming can indicate the body parts to be groomed (Gupta and Sinha, 2016), but the giver of grooming might also use subtle cues to dictate the interaction. There is no evidence for these possibilities at the neural level, since establishing this will require simultaneous brain recordings to study interbrain synchrony, and simultaneous tracking of both individuals’ joints to study how brain activity tracks joint movements of the self and partner.

Finally, social interactions require tracking of the grooming state as well as the social partner’s identity and movements. Which brain regions track these important social variables? While the grooming state (giving versus receiving) might be easily decoded through variations in sensory or motor activity during these states, precisely how each socially interacting individual keeps track of their partner’s identity or movements has not previously been demonstrated. For instance, social interactions often involve the two individuals not directly facing each other, such as when one monkey grooms the other’s back. While neurons in the high-level visual regions, such as the inferior temporal cortex, show invariant encoding of face identity (Tsao and Livingstone, 2008; Deen et al., 2023), but whether they maintain such representations actively throughout an extended social interaction is unclear. Likewise, the premotor cortex has mirror neurons which respond to movements made by an individual and movements that are observed (Rizzolatti and Craighero, 2004), but whether such mirror-neuron encoding of the partner’s joints is present during social interactions and whether it is modulated by grooming state is unclear. Addressing these questions will require recording brain activity not only during social interactions but also during more controlled cognitive tasks where basic visual and motor responses can be characterized.

Addressing the above fundamental questions also requires overcoming several methodological challenges. First, brain activity must be recorded wirelessly and simultaneously from socially interacting monkeys in a natural setting without any constraints. This requires not only seamless integration of two wireless recording systems but also ensuring social compatibility of the two monkeys to minimize the risk of aggressive encounters that could damage their implants. Second, since brain activity during social interactions is widespread, multiple high-level sensory and motor regions must be monitored simultaneously. Finally, for brain activity during social interactions to be more interpretable, it must be combined with brain recordings in a more controlled setting where basic sensory and motor responses can be characterized.

### Overview of the present study

Here, we addressed these issues through simultaneous wireless brain recordings from high-level visual and motor regions of two socially compatible monkeys while they interacted freely in a naturalistic setting without any constraints. We also recorded brain activity in each monkey separately while he performed visual and motor tasks on a touchscreen. We targeted the inferior temporal (IT) cortex, a region critical for object recognition that is also sensitive to social stimuli (Morin et al., 2015; Testard et al., 2024), the ventral premotor cortex (PMv), which contains mirror neurons sensitive to social gaze and intention (Rizzolatti and Craighero, 2004) and the ventrolateral prefrontal cortex (PFC), a region involved in executive control and social behaviors (Chang et al., 2013; Báez-Mendoza et al., 2021; Franch et al., 2024). Our results show that important social variables, such as a social favor signal, partner identity and partner movements are dynamically tracked throughout the social interaction, providing a rich neural basis for primate social behaviors.

## RESULTS

To investigate the neural basis of social behavior, we implanted two socially compatible male bonnet macaque monkeys (M1 and M2), each with 256 microelectrodes spanning high-level visual and motor regions (visual regions: inferior temporal cortex - IT; motor regions: ventral premotor cortex - PMv and ventrolateral prefrontal cortex – PFC; see Methods). We recorded brain activity wirelessly and simultaneously from both monkeys while they interacted freely without any constraints. In order to interpret brain activity as being of sensory or motor origin during the social interaction, we also recorded brain activity from each monkey separately while he performed a visual task and a motor task on a custom-designed touchscreen workstation with eye tracking capabilities (Jacob et al., 2021). In the visual task, each monkey had to fixate on face and body images of himself, his partner, and another unrelated monkey. In the motor task, each monkey had to touch targets appearing on a touchscreen with his left or right hand.

During the social interaction session, the two monkeys took turns grooming each other, interspersed with transition periods of no grooming (Figure 1A). Due to concerns about the two monkeys potentially damaging each other’s brain implants, we only recorded brain activity during a single brief social interaction (see Methods). Both monkeys exhibited similar reciprocal social interactions even before being implanted for brain recordings. Brain activity recorded during this entire session from the visual and motor regions showed rich dynamics (Figure 1B) that we did our best to elucidate in the sections below.

**Figure 1:**
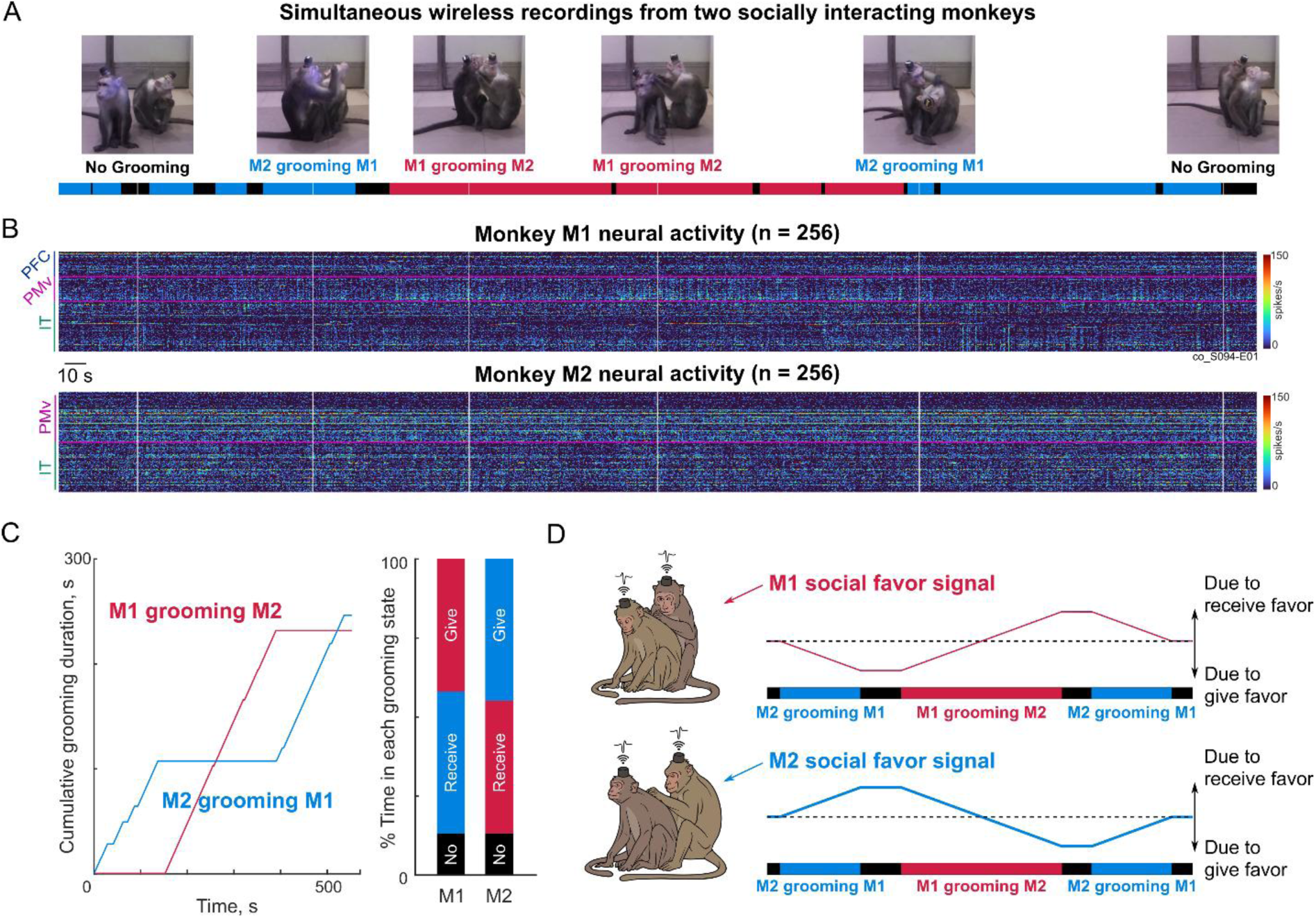
Simultaneous wireless brain recordings in socially interacting monkeys. (A) Timeline of the social interaction session with two monkeys, with example frames (*white lines*) illustrating the three types of grooming behavior observed: No grooming (*black*), M2 grooming M1 (*blue*) and M1 grooming M2 (*red*). (B) Firing rates in 0.5 s bins of all 512 neurons recorded wirelessly and simultaneously from both monkeys, with *white vertical lines* corresponding to the example frames in (A), and *pink horizontal lines* segregating brain regions (PFC, PMv, IT in M1; PMv, IT in M2). (C) *Left:* Cumulative duration of each type of grooming as a function of time: M2 grooming M1 (*blue*) and M1 grooming M2 (*red*). *Right:* Percentage of total session time spent by each monkey in grooming (marked *giving*), receiving grooming (marked *receiving*) and neither (marked *no*). Note that giving grooming for one monkey will be receiving grooming for the other, so the two bars are redundant. Both monkeys showed a near-perfect grooming reciprocity, i.e. groomed each other for almost equal durations. (D) Schematic of how monkeys’ brains might achieve reciprocity. Consider a session in which first M2 grooms M1, then M1 grooms M2 and then M2 grooms M1. We reasoned that each monkey could create a “social favor” signal representing the net duration spent in giving minus receiving grooming. This signal is positive if he has given more than he has received and negative if has received more than he has given. *Top:* For monkey M1, the social favor signal goes negative at first (since his partner is grooming him), then becomes positive and goes down to zero. *Bottom:* For monkey M2, the social favor signal goes positive initially (since he is grooming his partner), becomes negative and then rises back to zero.

We hypothesized that, during the social interaction, neural activity in each monkey should keep track of important social variables, such as the time spent in giving versus receiving grooming, the identity of the social partner and his movements. We also hypothesized that there should be dynamic interbrain coupling during the social interaction, which we sought to elucidate using predictive modeling.

### Social interaction & grooming reciprocity

The two male monkeys were socially compatible, with M2 dominant over M1. This dominance was also evident during the social interaction session in several ways: M2 mounted M1 at the start of the interaction, moved confidently around M1 brushing against him, while M1 remained stationary, M2 groomed M1’s face and handled M1’s implant to orient his head, and initiated grooming transitions. By contrast, M1 showed subordination to M2 in several ways, such as frequent lip-smacking, remaining stationary with his head down when M2 moved around him, grooming M2’s body more often than the face, and never handling M2’s implant. During the social interaction session, M2 groomed M1 first, M1 then groomed M2 and finally, M2 groomed M1 – all using stereotypical allogrooming behaviors (Figure 1A). While being groomed, both monkeys often used referential gestures to indicate the body part that they wished to be groomed, and their partner often responded immediately by grooming that body part ([Video S1](https://osf.io/fdjwa)). Such referential gestures have also been observed in the wild (Sinha, 1998; Gupta and Sinha, 2016).

Interestingly, M2 groomed M1 for a total of 244 s across 8 bouts, and M1 groomed M2 also for a similar total duration, of 230 s across 4 bouts (Figure 1C). Thus, both monkeys groomed each other for roughly equal durations (237 ± 7s), demonstrating an almost perfect reciprocity of grooming, in keeping with previous observations (Adiseshan et al., 2011; Onishi et al., 2013; Molesti and Majolo, 2017; Testard et al., 2024).

What could be the neural basis for this remarkable reciprocity? One possibility is that each monkey could create a running “social favor” signal, which tracks the net social favor (given minus received). In this case, social favor is quite simply, net duration of grooming (give – receive) or grooming surplus (Figure 1D) – but this could be more complex in the case of more sophisticated social interactions. A positive value of the signal indicates that the monkey has given more grooming than he has received. Presumably, when this signal reaches a high enough value, the monkey might decide to stop grooming and wait for his partner to reciprocate. Conversely, a negative value would mean that the monkey has received more grooming than he has given. Likewise, when the signal is sufficiently negative, the monkey might decide to stop receiving grooming and start grooming his partner. Critically, the grooming surplus signal will be opposite in sign for the two monkeys: when M1 is grooming M2, grooming surplus of M1 would increase whereas the grooming surplus of M2 would decrease. Further, we speculated that, since neural activity in motor regions would be directly involved in grooming behavior, such a social favor signal might be reflected in high-level visual regions (or other non-motor regions) rather than in motor regions, to avoid interference with any ongoing motor movements.

In our social interaction session, since at first M2 groomed M1, then M1 groomed M2 and then M2 groomed M1 again, the social favor (grooming surplus) signal for M1 would decrease at first, increase and then decrease again. Conversely, the social favor signal for M2 should increase at first, decrease and then increase again.

### Social favor signals as a neural basis for grooming reciprocity

To investigate whether the grooming surplus (social favor) signal in each monkey is represented in the population activity of each brain region, we plotted this signal along with the mean firing rate for each region. Since these quantities are likely to be slow varying signals, we plotted the firing rates and grooming surplus in 30 s bins. We obtained qualitatively similar results on varying this choice.

Strikingly, the average neural activity in IT of monkey M1 was strongly correlated with his own grooming surplus signal (*r* = 0.65, *p* < 0.005; Figure 2A, *upper panel*). Likewise, the average neural activity in IT of monkey M2 mirrored his own grooming surplus signal (*r =* 0.5*, p < 0.05*; Figure 2A, *lower panel*). This correlation was not present in the average neural activity of PMv neurons (correlation: r = 0.02, p = 0.94 for M1; r = 0.33, p = 0.17 for M2), nor in the average neural activity of PFC (r = 0.17, p = 0.5).

**Figure 2.**
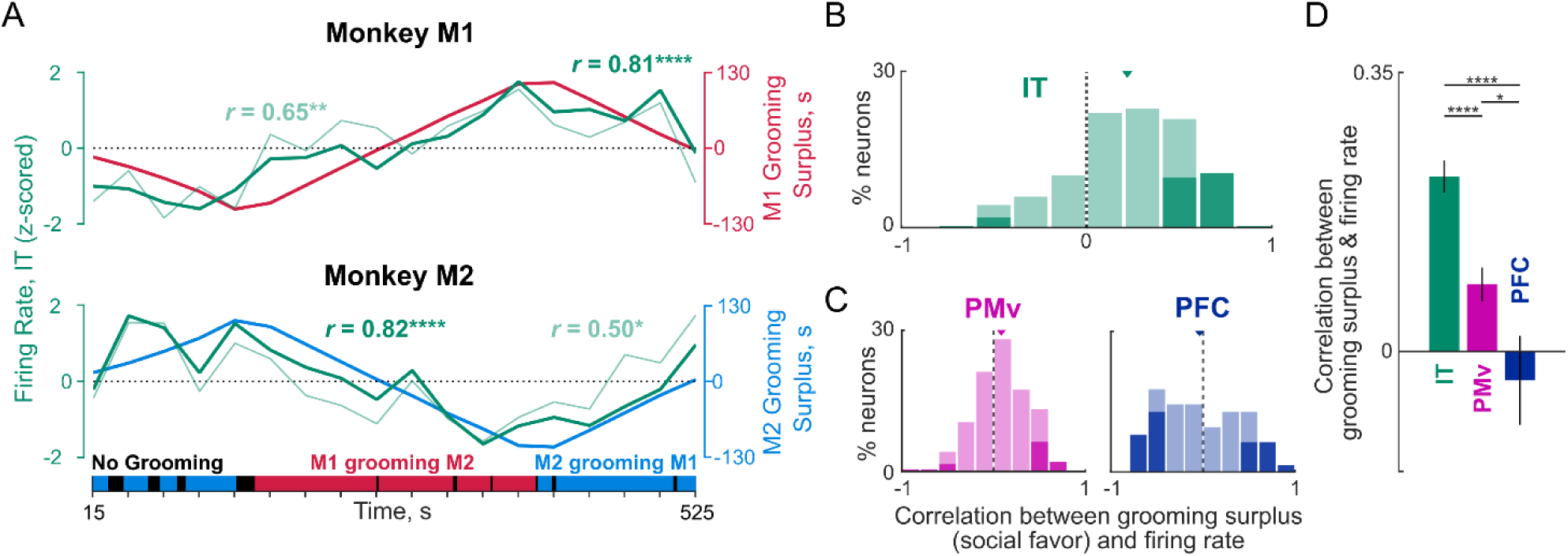
Reciprocity of grooming and its neural basis. (A) *Top:* Grooming surplus (social favor) signal for M1 (*red*), defined as the net duration of grooming (give – receive), overlaid with the average firing rate in 30 s time bins across all IT neurons in M1 (*thin green line*), and average firing rate across M1 IT neurons with a significant positive correlation with grooming surplus (*thick green line,* n = 28 out of 118 neurons). The number of such significant positive correlations were significantly different from the 2.5% expected by chance (p < 0.00005, chi-squared test). *Bottom:* Grooming surplus (social favor) signal for M2 (*blue*) overlaid with the average firing rate of all IT neurons in M2 (*thin green line*), and M2 IT neurons with a significant positive correlation (*thick green line*, n = 23 out of 125 neurons). The number of such significant positive correlations was also significantly different from the 2.5% expected by chance (p < 0.00005, chi-squared test). (B) Histogram of correlation between firing rates of neurons in each brain region with the respective grooming surplus (social favor) signal pooled across both monkeys for IT (*green*). Dark bars represent statistically significant correlations (p < 0.05), and the inverted triangle above each histogram represents the population mean. (C) Same as (B) but for PMv (*left panel, purple*) and PFC (*right panel, blue*). (D) Bar plot of average correlation (mean ± s.e.m across neurons, pooled across monkeys) with grooming surplus across neurons in each region. Asterisks indicate statistical significance of comparisons between brain regions (**** is p < 0.00005, * is p < 0.05, rank-sum test across correlation coefficients of neurons in the respective brain regions). For a comparison of whether grooming surplus is uniquely encoded compared to other alternate grooming-related variables, see Supplementary Section S2.

To investigate this further at the level of single neurons, we repeated this analysis on the firing rates of individual neurons. Across both monkeys, a substantial fraction of IT neurons (21%, 51 of 243 neurons) showed a significant positive correlation (p < 0.05) with the grooming surplus signal, while only 2% (6/243) showed a significant negative correlation (Figure 2B). In PMv, we observed fewer neurons (9% i.e. 16/185) with significant positive correlation and 3% (5/185) with significant negative correlation (Figure 2C, *left*). We found roughly equal proportions of neurons with positive and negative correlations with grooming surplus in PFC (positive correlation: 15% i.e. 9 of 61 neurons; negative correlation: 21% i.e., 13/61; Figure 2C, *right*). Thus, all three regions contained neurons that tracked the grooming surplus. Across the entire population of neurons, both IT and PMv neurons showed a significant positive average correlation with the grooming surplus signal (Figure 2D). By contrast, PFC neurons did not show a significant correlation with grooming surplus (Figure 2D). The grooming surplus correlation was strongest on average in IT, followed by PMv, then by PFC (Figure 2D).

### Unique encoding of grooming surplus compared to other variables

The above correlations with grooming surplus could arise from simpler differences in overall firing while giving and receiving grooming. Likewise, animals might keep track of grooming reciprocity by counting the number of times groomed, instead of the total duration. It is also possible that the observed correlation between neural responses and the groom surplus could be due to variations in the animals’ arousal or attentional states, as indexed by the alpha-band power. To resolve these issues, we created a number of possible alternative factors that could explain the temporal variation in firing rates observed in single neurons. For each factor, we calculated the average correlation and also its partial correlation with neural firing after removing all the other factors. This revealed that grooming surplus had the highest partial correlation with IT neurons even after removing all other factors including alpha-band oscillations (Section S2). Similarly, grooming state had the strongest influence on PMv neurons even after removing all other factors (Section S2). Interestingly, none of these factors were strongly encoded by PFC neurons (Section S2), but this null result is also limited by the fact that we only sampled PFC in one monkey.

We conclude that neural activity in IT closely tracks social favor (grooming surplus) signals in both monkeys. This unusual correspondence suggests that IT cortex helps maintain grooming reciprocity, or at the very least, reflects this signal after it is generated in other brain regions.

### Neural encoding of grooming state during social interaction

Social interactions involve not only tracking the relative duration of social favors like grooming, but also tracking of the ongoing grooming state. We therefore wondered whether the neural activity in the recorded brain regions encodes these grooming states. To start with, we plotted the average firing rate of each neuron when each monkey groomed his partner (i.e. giving grooming) against when it received grooming from his partner (i.e. receiving grooming). This revealed a heterogeneous pattern across neurons in each brain region (Figure 3A), with some neurons showing consistent increases, some showing decreases, and others showing no change in activity (Figure 3B).

**Figure 3:**
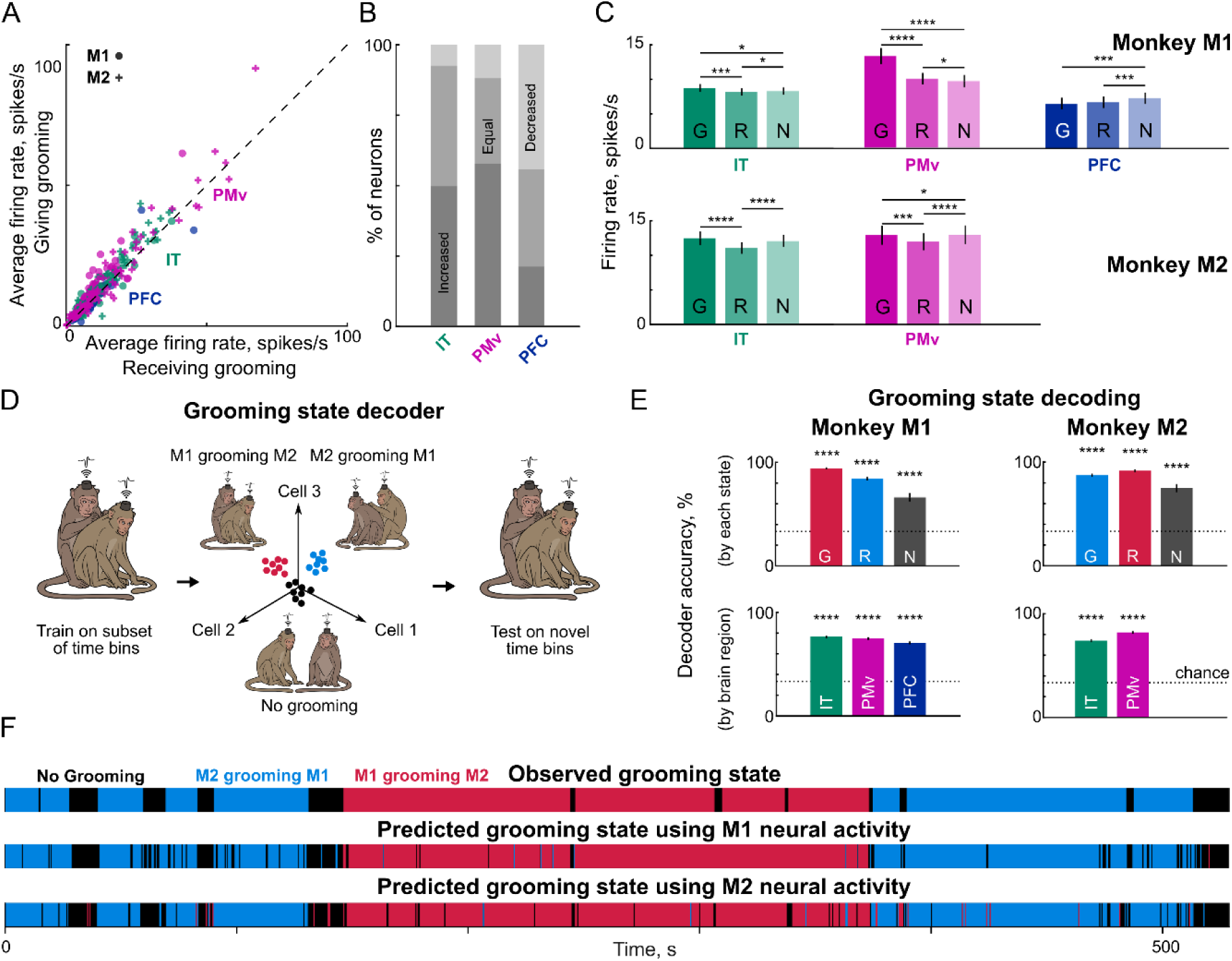
Neural tracking of grooming state during social interaction. (A) Average firing rate of all recorded neurons during Giving Grooming bouts plotted against the firing rate during Receiving Grooming bouts (*crosses*: M2 neurons; *dots*: M1 neurons; *green*: IT, *purple*: PMv, *blue*: PFC). (B) Percentage of neurons in each brain region whose firing showed a significant increase (*dark*), decrease (*light*) or no significant change (*medium*), as identified using a rank-sum test on their firing rate in 0.5 s bins across the social session (p <0.05). (C) Average firing rate in monkey M1 (*top*) and M2 (*bottom*) across neurons in each brain region during different grooming bouts: giving grooming (G), receiving grooming (R) and no grooming (N). Error bars indicate s.e.m across neurons. Asterisks indicate statistical significance of comparing mean firing rates across different grooming behaviors (* is p < 0.05, ** is p < 0.005, etc., sign rank test across mean firing rates of neurons during each grooming bout). (D) Schematic of grooming state decoding. We trained a linear classifier on the firing rates of neurons in 0.5 s bins labelled with their corresponding grooming state labels (M1 grooming M2, M2 grooming M1 or No grooming), and tested the classifier on novel time bins not seen during classifier training. (E) *Top panel:* Grooming state decoding accuracy for each monkey, of a classifier trained on all neurons pooled across brain regions for each grooming state: Giving (G), Receiving (R) and No Grooming (N). Accuracy was calculated by pooling the results obtained from 5-fold cross-validation across folds. Error bars represent s.e.m across 0.5 s time bins. Asterisks represent statistical significance of decoding accuracy (**** is p < 0.00005 using a chi-square test). *Bottom panel:* Grooming state decoding accuracy for each monkey, of a classifier trained on each brain region (IT, PMv, PFC) pooled across grooming states. Asterisks represent statistical significance calculated as before. (F) Observed grooming state for the entire session (*top*) shown along with predicted grooming state from the classifier trained on all neurons (across brain regions) from M1 (*middle*) and all neurons from M2 (*bottom*).

In both monkeys, the average firing rate in IT and PMv showed an increase in the giving grooming state compared to other conditions (Figure 3C). The increased PMv activity while grooming the partner presumably reflects the greater amount of movements involved during these bouts. Indeed, we found that the neurons that showed increased activity during the touchscreen-based motor task also showed greater activity while giving compared to receiving grooming (Section S1). Likewise, the increased IT activity during giving grooming could be due to increased visual inputs during giving grooming. Indeed, we found that neurons that showed increased activity during a fixation task showed a larger difference in activity during giving vs receiving grooming (Section S1).

The increased firing rates during giving compared to receiving grooming is interesting, but to be expected since there is greater sensory and motor engagement for the animal when it is grooming his partner. However, we observed an opposite pattern in the local field potentials: there was greater LFP power in both IT and PMv while receiving grooming compared to giving grooming in both monkeys (Section S3). We also observed greater LFP power particularly in the alpha band in IT cortex during receiving grooming (Section S3), presumably due to the monkey closing his eyes or relaxing while receiving grooming. We speculate that the LFP activity could reflect greater inhibitory inputs while receiving grooming, which in turn results in reduced neural firing.

Since neural responses are systematically different across grooming states, we wondered whether grooming state can be reliably decoded from the neural activity recorded during the social interaction. To this end, we trained a linear classifier on the firing rates of neurons in successive 0.5 s time bins throughout the social session (Figure 3D). To avoid overfitting we used a 5-fold cross-validation procedure where the grooming state in a given time bin was always predicted using a classifier that was never trained on this bin. Decoding accuracy was consistently above chance for all grooming states (Figure 3E) and in each brain region in both monkeys (Figure 3E).

To visualize the dynamics of grooming state decoding, we plotted the observed grooming state throughout the session and compared it against the grooming state predicted by the neural decoder of each monkey. This revealed a striking match between the observed and predicted grooming state in both monkeys (Figure 3F). Finally, we wondered whether the decoders trained on grooming states could predict the onset of the grooming state using neural activity even before it was evident in behavior. This was indeed the case: the decoder probability began to rise a few seconds before the onset of giving or receiving grooming (Section S4).

Thus, each grooming state during the social interaction elicited distinct patterns of neural activity in both visual and motor regions.

### Neural tracking of partner identity during social interaction

Social interactions require not only tracking social favors and grooming state, but also the identity of the current social partner. We therefore wondered whether neural activity in both monkeys keep track of the identity of the partner during the social interaction. Since neural activity during social interaction involves a multitude of signals, we can only address this question by recording neural activity in a more controlled setting. To this end, we recorded neural responses while each monkey performed a fixation task (before the social session), where he viewed face & body images of himself, of his partner and of a third unrelated monkey (Figure 4A). We then trained 3-way linear decoders to decode individual monkey identity from the population activity recorded in each region and tested them in the social session (Figure 4B).

**Figure 4:**
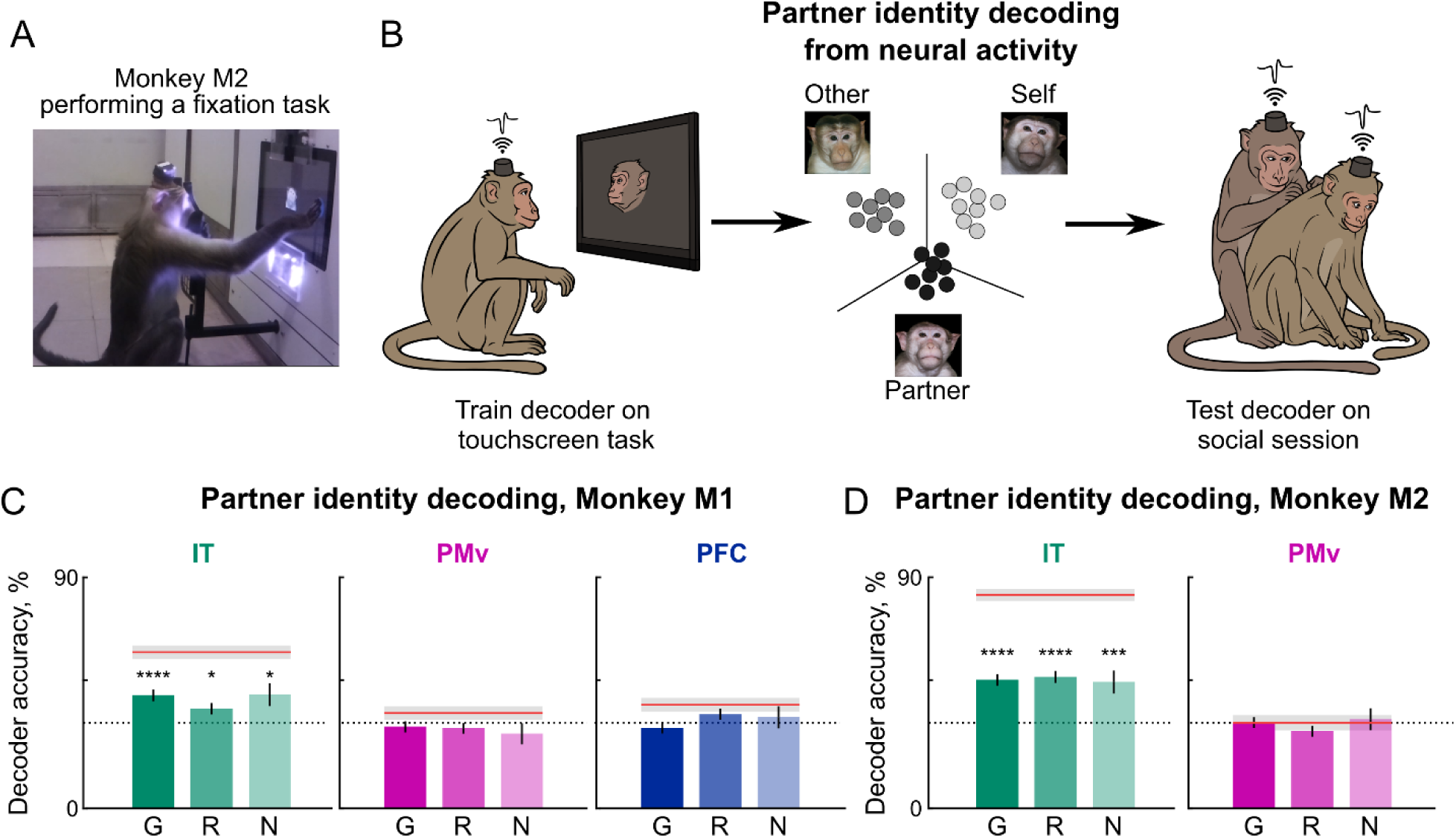
Neural tracking of partner identity during social interaction. (A) Example frame showing monkey M2 performing a fixation task. (B) To investigate if each monkey’s brain tracks his partner identity during the social interaction, we recorded neural responses while the monkey viewed face photographs of himself, his partner and a third monkey on a touchscreen (*left panel*). We then trained a linear decoder to decode face identity (*middle panel*), and tested this decoder on neural activity during the social interaction (*right panel*). (C) Partner identity decoding accuracy in IT, PMv and PFC neurons of Monkey M1 during the social interaction session, separated into the three grooming bouts: giving grooming (G), receiving grooming (R) and no grooming (N). Decoding accuracy was calculated in 0.5 s bins. Error bars represent s.e.m across all bins of that type. Decoding accuracy in the touchscreen task is also shown for reference (*red lines*, with *shaded regions* indicating s.e.m). Asterisks represent statistically significant decoding accuracy (* is p <0.05, **** is p < 0.00005, chi- square test of correct and incorrect counts across all bins compared to the numbers expected by chance). (D) Same as (C) but from the brain regions in Monkey M2.

In Monkey M1, neural activity in IT (but not PMv & PFC) correctly decoded his partner i.e. Monkey M2 in all grooming states (Figure 4C), but particularly so while giving grooming & no grooming, and particularly so when receiving grooming on his face from his partner (Section S5). We speculate that this could be due to Monkey M1 tracking his partner more closely, since he was the subordinate. We observed similar results in Monkey M2: neural activity in IT (but not PMv) correctly decoded his partner i.e. Monkey M1 across all grooming states (Figure 4D), with no obvious differences in decoding across states (giving/receiving groom) or across body parts (Section S5). Partner decoding changed dynamically over time in general (Section S5), sometimes in ways related to the ongoing behavior (Section S9). We were also able to decode the hand being used by each monkey during the social interaction, based on using PMv responses in the motor task (Section S5).

Thus, neural activity in IT tracks the social partner consistently throughout the social interaction. Interestingly, we did not observe differences in partner decoding, even in grooming states where the monkey may not have been directly looking at his partner.

### Neural tracking of self and partner movement during social interaction

Social interactions inevitably involve closely coordinating one’s own movements with the partner depending on the nature of the ongoing interaction. We therefore wondered whether neural activity in each monkey would track his own and his partner’s movements throughout the social interaction, and whether this would be dependent on the nature of grooming interaction.

We had recorded videos from multiple views of the entire social interaction that were synchronized to the wireless neural recordings. We selected the top view videos for further analysis since most joints of both monkeys were clearly visible without occlusions (Figure 5A). Since existing automated pose tracking algorithms did not work well to separately identify the joints of the two interacting monkeys, we identified 20 key joints of each monkey for manual annotation, and got them manually annotated and cross-verified by a team of annotators (Figure 5A). We used the head, neck, left and right shoulder positions as regressors, since these joints were visible in a large number of frames (we obtained qualitatively similar results on varying this choice). We then built a model for each neuron where the firing rate in successive time bins was predicted as a linear combination of the monkey’s own joint positions, or his partner’s joint positions (Figure 5A; see Methods).

**Figure 5.**
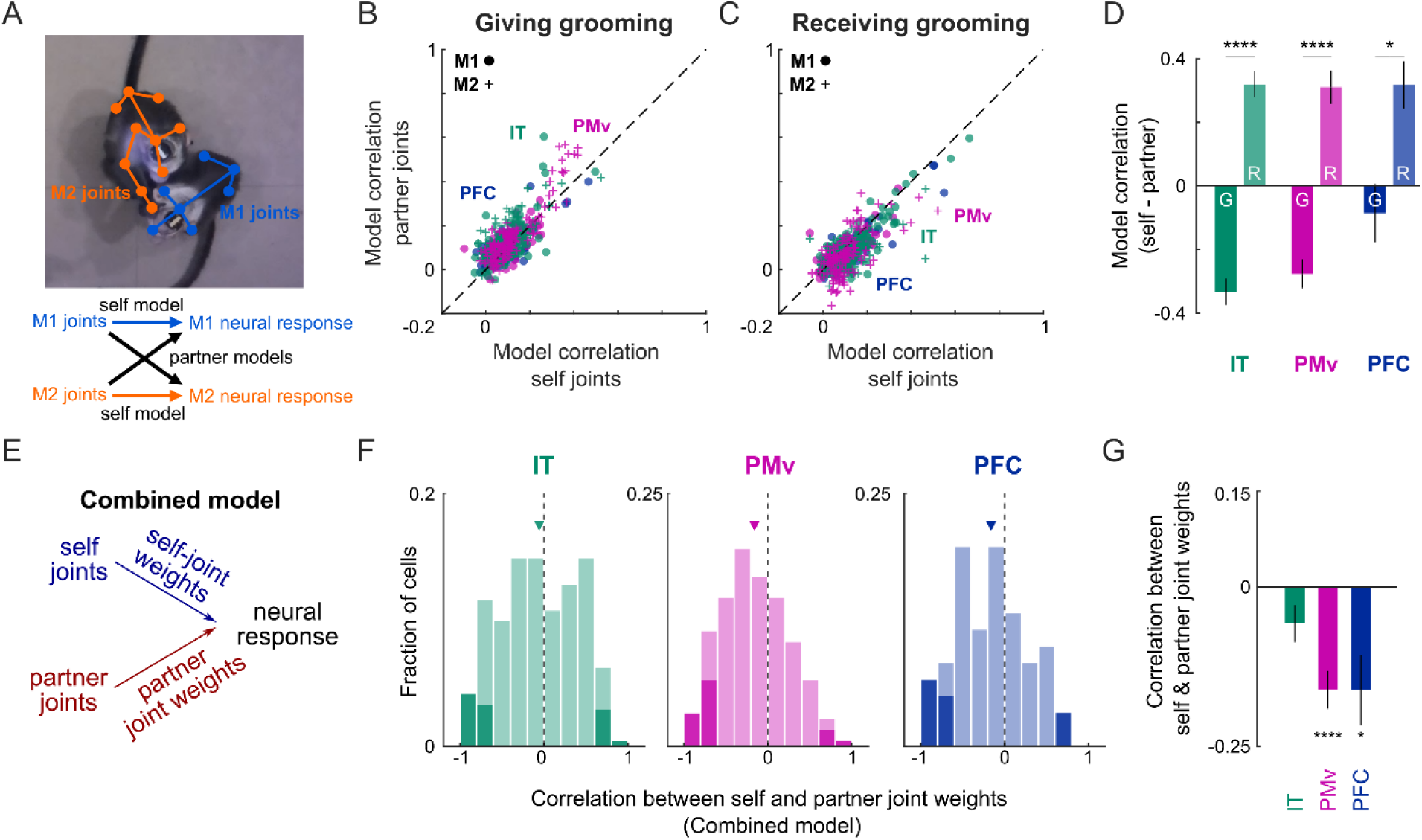
Neural tracking of self and partner movements during social interaction. (A) *Top:* Example screenshot from the top view of the two socially interacting monkeys, with manual joint annotations marked for M1 (*blue*) and M2 (*orange*). *Bottom:* schematic of joint models: we predicted the firing rate of each neuron in M1 and M2 using the manually annotated joint positions of M1 and M2. (B) Cross-validated correlation between predicted and observed firing rate across time bins of the partner joint model plotted against that of the self-joint model for each neuron in IT (*green*), PMv (*purple*) and PFC (*blue*) for monkeys M1 (*dots*) and M2 (*crosses*) during the giving grooming condition. Here, the partner joint model performed better than the self joint model. (C) Same as (B) but in the receiving grooming condition. Here, the self-joint model performed better than the partner joint model. (D) Difference in correlation between self and partner joint model correlations (mean ± s.e.m) for IT (*green*), PMv (*purple*) and PFC (*blue*) in the giving (G) and receiving\ (R) grooming conditions. Asterisks indicate significance of the model correlation comparison, obtained using sign-rank test (* is p < 0.05, **** is p < 0.00005). (E) Schematic of the combined model in which the firing rate of each neuron in successive time bins was predicted as a linear combination of self and partner joints. (F) Histogram of correlation between self and partner joint weights across neurons in IT (left), PMv (middle) and PFC (right), with inverted triangles indicating the means. Dark bars indicate neurons with significant correlation (p < 0.05) (G) Correlation (mean ± s.e.m) between the model weights for the self joints and partner joints for IT (*green*), PMv (*purple*) and PFC (*blue*). Asterisks indicate statistically significant deviations from zero, obtained using sign rank test (* is p < 0.05, **** is p < 0.00005).

We then plotted the cross-validated model correlation of the partner joints model for each neuron against that of the self joints model for giving grooming (Figure 5B) and receiving grooming (Figure 5C). We observed a striking dissociation: in all brain regions, neural responses of the groomer were always predicted better by the receiver’s joints during giving grooming in both monkeys (Figure 5D). Conversely, while receiving grooming, neural responses of both monkeys were predicted better by their own joints than by the joints of their partner who was grooming them (Figure 5D). This pattern unfolded dynamically over time, revealing rises and falls presumably at times when the animal was more closely tracking his partner (Section S6).

Thus, neural activity in both the groomer’s and receiver’s brains are strongly driven by the grooming receiver’s joint positions, suggesting that the monkey receiving groom drives the ongoing interaction.

### Neural mirroring of joint position tuning during social interaction

Social interactions require tracking our partner’s intentions and movements and linking them to ours. Indeed, “mirror neurons” have been reported in premotor regions which have congruent tuning for self and partner movements (Rizzolatti and Craighero, 2004). To investigate this issue further, for each neuron, we built a model where its firing rate was predicted as a linear combination of both the self joint positions and partner joint positions (Figure 5E).

In this combined model, we calculated the correlation between the model weights for self joints and partner joints. A positive correlation would imply that single neurons have similar tuning for self and partner joints, as might be expected for mirror neurons. A lack of correlation (i.e., zero correlation) would imply that different populations of neurons keep track of self and partner joints. A negative correlation would imply that observing a partner moving his joint would reduce the firing rate in neurons coding for that joint movement in oneself, which could be potentially useful to avoid making self movements while observing others’ actions. All three possibilities are conceptually interesting, and have not been studied systematically during natural social interactions in primates.

In all three brain regions but most prominently in PMv, we observed a negative correlation on average between model weights for self and partner joints, consistent with the third possibility above (Figure 5G). At the single neuron level, however, there was considerable diversity, with neurons showing both significant positive and negative correlations (Figure 5F; positive correlations: 3.3 % of 243 neurons in IT, 2.2% of 185 in PMv, 3.3% of 61 in PFC; negative correlations: 7.4 % in IT, 9.7% in PMv, 11.5% in PFC). Although relatively few neurons showed significant correlations, these fractions are substantially different from the 5% expected purely by chance.

We conclude that a subset of neurons in high-level sensory and motor regions show congruent encoding of joint positions of both self and partner joints, providing a neural basis for tracking the intentions of the social partner.

### Interbrain coupling during social interaction

Social interactions can involve varying degrees of coordination between the interacting partners, which might result in varying inter-brain coupling between the interacting brains. In this particular dyadic interaction, the two monkeys were either giving or receiving groom, and we wondered whether their brains were coupled differently during these two grooming states. To investigate this issue, in each monkey, we built a linear model to predict the activity of each of his neurons using the population activity of his partner (Figure 6B, see Methods).

**Figure 6.**
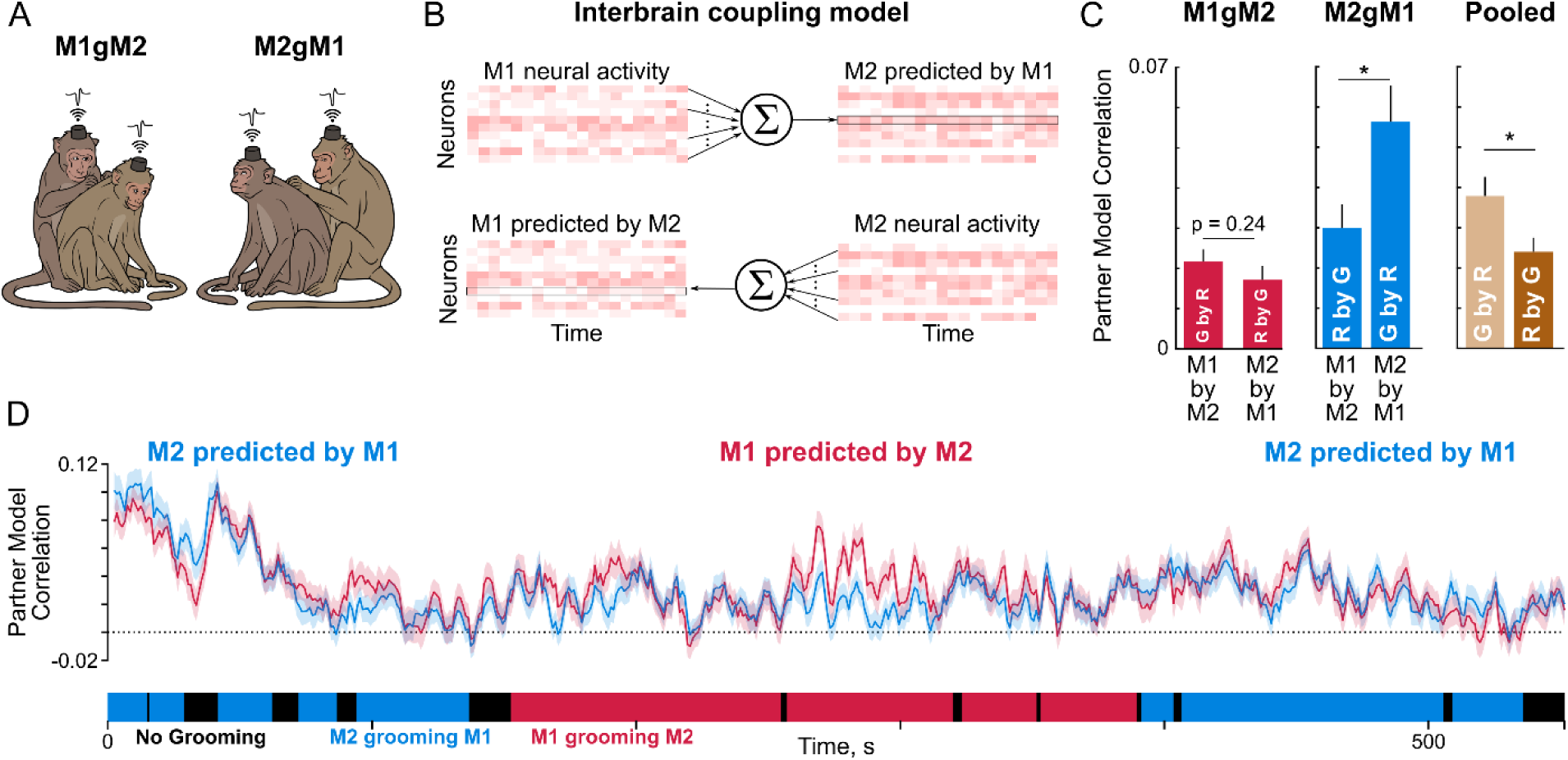
Interbrain coupling and its modulation by grooming state in PMv/PFC. (A) Schematic of the two grooming states observed during social interaction: M1 grooming M2 (*left*) and M2 grooming M1 (*right*). (B) Schematic of the interbrain coupling model. For each monkey (M1/M2), we built a model to predict the firing rate of his neurons using the PMv and PFC population neural activity of his partner, in 50 ms time bins (see Methods). (C) Partner model correlation (mean + s.e.m) for M1 and M2 while M1 groomed M2 (*left, red bars*), while M2 groomed M1 (*middle, blue bars*), and for the data pooled for the groomer and receiver monkeys (*right, G by R* is the partner model correlation for the groomer’s neurons predicted by the receiver’s neurons; *R by G* is partner model correlation for the receiver monkey neurons). Asterisks indicate statistically significant comparisons (* is p < 0.05, rank-sum test across neurons). (D) Time course of smoothed partner model correlation during the entire interaction (mean ± s.e.m across 50 ms time bins in each 5 s window). It can be seen that partner model correlations vary dynamically across the entire social interaction.

In both monkeys, there was a substantial fraction of PMv/PFC neurons whose activity was significantly predicted by the partner’s brain activity (64% in M1, 52% in M2, *p* < 0.05 for partner model correlation). To investigate whether this interbrain coupling differs across grooming states, we calculated the average partner model correlation for the significantly modulated M1 and M2 neurons for each type of grooming. This revealed an interesting double dissociation: the groomer’s brain activity was consistently predicted better by the receiver compared to vice-versa (Figure 6C).

The relatively small partner model correlations in this analysis could arise from the fact that we are predicting the firing rates in small time bins for each channel in a single long trial. To evaluate this possibility, we calculated for each channel the average firing rate profile, and generated two Poisson spike trains from this firing rate profile, and calculated the correlation between their firing rates. This yielded relatively small correlations (correlation, mean ± s.e.m across channels: *r* = 0.055 ± 0.005 for M1, 0.045 ± 0.004 for M2). Thus, the relatively small magnitude of partner model correlations are due to the low correlation expected from single-trial firing rates.

Since social interactions are dynamic, we wondered whether interbrain coupling would unfold dynamically over the course of the social interaction. To this end, we took the instantaneous correlation at each 0.05 s time bin between the predicted and observed firing rates across all neurons, and smoothed this instantaneous correlation with a window of 5 s which matched the slower movement timescales present in behavior. This revealed a dynamic picture with larger interbrain coupling near transition points between grooming states as well as within each grooming state (Figure 6D). We observed similar interbrain coupling patterns for the other brain regions as well (Section S7). We also looked for but did not find systematic differences in temporal delays or coherence at specific frequencies as a function of grooming state (Section S8).

We conclude that social interactions involve dynamic interbrain coupling that is driven by the receiver of grooming rather than the groomer. This finding is consistent with our earlier observation that the groomer’s neural activity is driven by the receiver’s joints (Figure 5). Indeed, we observed several occasions during the social interaction where the receiver adjusted his position to indicate where he wanted to be groomed, demonstrating that the social interaction was indeed driven by the receiver and not the groomer ([Video S1](https://osf.io/fdjwa)).

### Convergence between behavior and multiple neural signals

In the preceding sections we have shown that, during the social interaction, brain activity in both monkeys revealed dynamically varying signals related to the grooming surplus, partner identity, partner joints, interbrain coupling and brain oscillations. Although we have reported that each signal was separately modulated by the grooming state and that each signal varied dynamically throughout the session, our analyses so far do not integrate the dynamics of these signals to interpret them in the light of the ongoing behavior. For instance, there could be specific behavioral events at which all signals show clear and interpretable modulation. Conversely, there could be times at which a neural signal attains a peak, and there is some interesting ongoing behavior at that time. To this end, we closely scrutinized the behavior of both monkeys as well as the corresponding neural signals, to find convergent and interpretable links. We summarize our observations briefly below, and refer the reader to Section S9 and [Video S2](https://osf.io/nfpr5) for details.

We noted several interesting convergences between behavior and all the neural signals. First, when M2 stopped grooming M1 (Video time = 266 s), M2 turned around and presented his back to M1, M1 huddled up against M2 and both made lip-smacking gestures and M1 began to groom M2. During this time, we found that M1 initially tracked his partner’s identity (when M2 stopped grooming M1), then M2 tracked his partner’s identity (when M1 began to groom M2), as evidenced by partner identity decoding. M1 closely tracked his partner’s joints, while M2 tracked both his own and partner’s joint movements. M1’s brain activity is better predicted by M2, indicating interbrain coupling. All these neural signals point to a consistent, ongoing internal state in the two monkeys.

Likewise, when M1 was grooming M2 (Video time = 520 s), M2 started grooming M1 and M1 immediately responded by stopping his grooming. During this time, M1 was relaxed, as evidenced by high alpha power. M2 was tracking his partner’s identity, as evidenced by partner decoding. Both monkeys were tracking the receiver’s joints. M2’s brain was predicted better by M1’s brain, indicating interbrain coupling.

Finally, there were several neural events where alpha power reached a peak in each monkeys’ brain. This was usually when that monkey was receiving grooming and looking up either at the roof or had his eyes closed. These observations suggest that the receiver of grooming is in a relaxed state. We also observed several moments when the alpha power attained smaller peaks while giving grooming, thus suggesting that the groomer also might be in a relaxed state while grooming his partner. This is consistent with similar observations made in the wild (Yates et al., 2022).

## GENERAL DISCUSSION

Here we investigated the neural basis of primate social interactions by simultaneous wireless brain recordings from the high-level visual and motor regions of two male bonnet macaque monkeys interacting naturally without any constraints. Our main findings are: (1) single neurons, primarily in IT, encoded a social favor signal representing the net duration spent in giving minus receiving grooming in both monkeys, providing a neural basis for grooming reciprocity; (2) neural activity in each monkey tracked the grooming state as well as the partner’s identity; (3) neural activity in both monkeys were driven strongly by the joints of the grooming receiver rather than the groomer; (4) neural activity in the groomer’s brain was predicted better by the receiver’s brain than vice-versa. These findings suggest that the monkey receiving grooming directed his partner’s interaction, which was also evident in behavior ([Video S1](https://osf.io/fdjwa)). Below we discuss these findings in relation to the existing literature.

### Neural encoding of social favors

A striking observation from our study is that neural activity in IT cortex of each monkey tracked the relative time spent in giving versus receiving grooming (Figure 2). This finding is congruent with the recent observation that IT and PFC neurons in individual monkeys interacting with a partner can predict net grooming duration (Testard et al., 2024). However, our study goes a step further to demonstrate a simultaneous correlation between each monkeys’ neural activity and his ongoing grooming surplus (Figure 2). We have also shown that grooming surplus was encoded most strongly by IT compared to PMv and PFC. At first glance, it is somewhat surprising that a visual region (IT) rather than motor regions (PMv/PFC) should encode the grooming surplus. However, we propose that this is quite reasonable considering that encoding such a signal, if encoded in the motor cortex, would interfere with ongoing motor output. We also caution that it is unlikely that IT would be the sole carrier of such a signal, since social interactions can evoke widespread brain activation and any of these brain regions could encode social favor signals (Wittmann et al., 2018; Freiwald, 2020; Deen et al., 2023).

Our finding that IT neurons encode a grooming surplus signal must also be carefully weighed against several alternative explanations, since giving and receiving grooming involve completely different visual inputs and motor actions. To rule out these possibilities, we constructed a number of alternative time-varying signals that could be encoded in the firing rate: these included binary signals that signal giving or receiving grooming, as well as other continuous signals that involve cumulative duration of giving or receiving grooming alone. Neural responses in IT remained significantly correlated with the groom surplus signal even after removing the contributions of these other signals (Section S2).

Another possibility is that the observed rise and fall of neural responses in each monkey signals a slow-varying arousal signal rather than a social favor signal. We consider this possibility unlikely. This is because arousal would typically be largest at the time of transitions between grooming states in both monkeys, which would give rise to a positive correlation between the firing rates in both monkeys with arousal – by contrast, their firing rates are in fact, negatively correlated. Nonetheless, to investigate this possibility further, we asked whether the correlation between neural responses and grooming surplus remained significant after regressing out the LFP alpha band power, which is widely taken as a measure of arousal or attentional state (Klimesch, 2012; Morrow et al., 2023). Thus, changes in arousal cannot explain the observed correlation between the groom surplus signal and neural responses.

### Encoding of grooming variables during social interactions

Previously, in monkeys trained to play social games with a partner under artificial constraints, prefrontal neurons keep track of the partner’s choices, reward states and strategies (Yoshida et al., 2012; Haroush and Williams, 2015; Ong et al., 2021; Chang et al., 2013; Báez-Mendoza et al., 2021). Our finding that neural activity tracks key variables like grooming state (Figure 3), partner identity (Figure 4) and partner’s joint movements (Figure 5) is congruent with these studies. However, our findings go a step further to demonstrate that such tracking occurs spontaneously in natural social interactions in monkeys without any extensive social training.

Our finding that grooming state can be decoded from neural activity in all regions (Figure 3) is to be expected, since the sensory inputs and motor outputs during each grooming state are likely to be quite different in nature. Perhaps what is more interesting is that grooming state could be continuously decoded throughout the social interaction (Figure 3F), even though sensory inputs or motor outputs were dynamically changing throughout.

Our finding that IT neurons track partner identity during the social interaction (Figure 4) is likewise expected given that they carry invariant information about objects and faces (DiCarlo et al., 2012). However, our finding of robust generalization of partner identity decoding from images on a screen to a real-world social interaction has not been reported previously. More interestingly, we have found that partner identity decoding was remarkably stable throughout, even while receiving grooming, when the monkeys were often looking away from their partner. Thus, IT neurons track socially relevant stimuli even when they are not directly visible. These findings are broadly consistent with the encoding of unseen stimuli in IT (Puneeth and Arun, 2016) as well as the encoding of social stimuli in the primate temporal lobe (Sliwa and Freiwald, 2017; Shepherd and Freiwald, 2018; Freiwald, 2020; Deen et al., 2023).

### Encoding of self and partner joints during social interactions

We have found that neural activity in all three recorded brain regions (IT, PMv, PFC) is reliably predicted by both self and the partner’s joint movements (Figure 5). Interestingly, in all three regions, brain activity was better predicted by the partner’s joints while giving grooming than while receiving grooming, and was better predicted by its own joints while receiving grooming.

We speculate that this could be because the receiver is driving the grooming interaction by constantly adjusting his position, causing the groomer to keep track of the receiver’s joints. We note that all three regions are well-positioned for this kind of tracking: IT could encode specific visual features of each joint, while PMv and PFC are involved in motor movements and also receive projections from IT, and could therefore encode both self and partner joints (Rizzolatti and Craighero, 2004). Interestingly, we found neurons with both positive and negative correlations in tuning for self and partner joints, which provide a neural basis for understanding the social partner’s intentions (Rizzolatti and Craighero, 2004). However, the prevalence of negative correlations between self and partner joint tuning suggests that in social interactions, a movement made by the social partner might have the overall effect of inhibiting a similar movement in oneself, while facilitating reciprocating joint movements. We speculate that this inhibition of joint-related activity could also provide a neural substrate for tracking partner joints, since the reduced neural activity could be used to interpret the social partner’s actions. Taken together, our study demonstrates the spontaneous tracking of partner joints during natural social interactions.

### Interbrain coupling during social interactions

We have observed substantial interbrain coupling that varies dynamically throughout the social interaction (Figure 6). This is consistent with previous reports of interbrain coupling in bats (Zhang and Yartsev, 2019; Rose et al., 2021), mice (Kingsbury et al., 2019; Zhang et al., 2025) and even humans (Stephens et al., 2010; Kinreich et al., 2017; Schilbach and Redcay, 2025). Such interbrain coupling during social interaction is presumably due to common sensory and motor events experienced by the interacting individuals (Schilbach and Redcay, 2025). However, interbrain coupling between socially interacting monkeys has never been observed previously due to the challenges involved in obtaining such brain recordings.

Our finding that the groomer’s brain activity is driven by the receivers’ brain activity (Figure 6) is consistent with our observation that the receiver’s joints drive the groomer’s brain activity (Figure 5) – these observations suggest that it is the receiver of grooming that drives the social interaction, and not the groomer. This finding is surprising at first glance, because one would expect the groomer’s movements to cause sensations in the receiver. However, a closer inspection of the social interactions revealed that the receiver monkey kept indicating where it wished to be groomed, suggesting that it indeed drove the social interaction ([Video S1](https://osf.io/fdjwa)). These referential gestures involve changes in body posture or position, which might be reflected in the receiver monkey’s brain activity. As the groomer monkey responds to such referential gestures and starts grooming at the indicated location, this might change the groomer monkey’s neural activity, resulting in the inter-brain coupling we see in this study. Such referential gesturing has also been observed in the wild (Gupta and Sinha, 2016). More broadly, our results show that interbrain coupling occurs dynamically throughout the social interaction but with directionality depending on the social context.

### Understanding social interactions in the wild

Our study represents a first step towards studying the neural basis of primate social interactions in a naturalistic setting. We have characterized brain activity during only one specific dyadic social interaction, but primate social behaviors are far more complex. For instance, allogrooming reciprocity could occur not just within a single session but can potentially take place over multiple interactions or sometimes not at all (Sinha, 1998; Adiseshan et al., 2011; Onishi et al., 2013; Gupta and Sinha, 2016; Molesti and Majolo, 2017). Non-reciprocal interactions frequently occur for other reasons as well. For instance, a subordinate might groom a dominant individual after an aggressive interaction, and this interaction is often one-sided. Likewise, subordinates or dominant individuals may initiate grooming towards another individual multiple times, but each interaction may be very brief and non-reciprocal (Ram et al., 2003; Sinha et al., 2005; Xia et al., 2013). We speculate that, in such cases, the social favor signal might start from a non-zero (positive or negative) value and go to a baseline level after the interaction is over. We also speculate that, since it could be tedious to maintain a non-zero social favor signal at the neural level, social interactions would be reciprocal by default unless there is some compelling reason for them not to be so.

Monkeys keenly observe other dyadic interactions in their social group to learn or update their knowledge of social dominance in their social group (Sinha, 1998, 2017). Interestingly, their behavior is predicted better by the duration rather than the frequency of grooming interactions between two individuals (Sinha, 1998). This is consistent with our observation that neural responses track the duration of grooming rather than the number of grooming bouts (Section S2). However, an individual might make many attempts to groom or solicit grooming directed at specific group members. We speculate that, in this case, neural activity might keep track of the frequency of these interactions as well. Indeed, we have found the number of grooming bouts also uniquely explains neural activity (Section S2). Based on all these observations, we speculate that the social brain of an individual maintains a long-term social ledger of his own interactions with each individual in their social group, as well as a record of the prevailing hierarchy of the entire social group (Sinha, 1998, 2017).

### Limitations of this study

Our study only represents a first step towards understanding social interactions between monkeys in a naturalistic setting, and has several limitations. First, we were only able to record simultaneous brain activity during a single brief social interaction, due to technical challenges involved in simultaneous recordings, and due to our concerns that both monkeys might damage each other’s sensitive brain implants. As a result our findings regarding the encoding of key grooming variables in the high-level visual and motor regions might not hold across many more social interaction sessions. We consider this highly unlikely, since the social interaction recorded in our study was representative of the many social interactions we have observed between the two monkeys in our study. The broader concern however is that social interactions may be more heterogeneous across sessions or more likely across multiple dyads of interacting individuals, and that our results may not hold across the wider range of possible social interactions. Whether our results generalize or not to the broader range of social interaction is an interesting possibility for future work. However, none of these concerns invalidate the robust tracking of grooming variables observed during the dyadic social interaction studied here.

Second, we have only recorded from a very limited set of brain regions in our study. This is an important limitation because social interactions evoke widespread neural activity throughout the brain (Sliwa and Freiwald, 2017; Shepherd and Freiwald, 2018; Freiwald, 2020; Deen et al., 2023). Thus the observed encoding of grooming variables in the high-level visual and motor regions might only reflect the encoding in other regions that may directly drive social interactions. Likewise other brain regions might encode other interesting social variables that were not observed in our study. Similarly, it could be that other regions in both monkeys’ brains, such as primary motor cortex, directly drive the monkeys’ own actions and do not show modulations with grooming state as we have observed here. The limited monitoring of only a few brain regions is a major limitation for any study that aims to understand primate social interactions, but these are challenges unique to primate neurophysiology in general. Addressing these limitations will require monitoring the whole brain with even more dense electrode arrays or using other methods or species where whole-brain monitoring is technically more feasible. These are exciting possibilities for future work.

### Conclusions

In sum, our results show that important social variables, such as the ongoing social favors, grooming state, partner identity and joint movements are encoded in high-level visual and motor regions, providing a rich and dynamic neural substrate for the ongoing social interaction.

## METHODS

All animal experiments were performed according to a protocol approved by the Institutional Animal Ethics Committee of the Indian Institute of Science, Bangalore (CAF/Ethics/750/2020) and by the Committee for the Purpose of Control and Supervision of Experiments on Animals, Government of India (V-11011(3)/15/2020-CPCSEA-DADF). Many experimental procedures are common to previously reported studies from our laboratory (Ratan Murty and Arun, 2015; Jacob et al., 2021), and summarized briefly below.

### Neurophysiology

We implanted two adult bonnet macaque monkeys (*Macaca radiata,* M1: *Co* & M2: *Di*, aged 9.5 years) with Floating Microelectrode Arrays (FMA) from Microprobes for Life Science using standard neurophysiological procedures. Each FMA had 32 single-shank microelectrodes arranged in a 4 x 8 grid with an inter-electrode spacing of 400 microns. Monkey M1 was implanted with 4 FMAs into the central portion of the left inferior temporal (IT) cortex, 2 FMAs in the left ventrolateral prefrontal cortex (PFC), and 2 FMAs in the left ventral premotor cortex (PMv). Monkey M2 was implanted with 4 FMAs in the right inferior temporal cortex and 4 FMAs in the right ventral premotor cortex. In both monkeys, these FMAs were connected to a custom-designed titanium pedestal fixed to the skull, containing headstages and wireless interface boards,. Each monkey underwent a single surgery under anaesthesia using standard procedures (Ratan Murty and Arun, 2015).

On the day of recording, each monkey was brought into a custom chair and its snout was restrained temporarily (for ∼5 minutes) in order to fix the wireless logger to the titanium pedestal using screws. Extracellular wideband signals were logged at 25 KHz using a custom wireless logger that was synchronized to a neural data acquisition system (*eCube,* White-Matter LLC). These wideband signals were high-pass filtered and thresholded using a commercial software (*OfflineSorter*, Plexon Inc) to obtain multiunit activity.

### Social interaction

The social interaction was conducted in a naturalistic environment described previously (Jacob et al., 2021). Five video cameras were positioned around the room to capture the behavior of the interacting animals from multiple views, of which three cameras were time-synchronized to the neural recordings by the neural data acquisition system, and the other two were synchronized offline using alternative methods.

Both monkeys were housed together in a larger social group since birth and so were already socially compatible. After including them into our study, we pair housed them fully until they were implanted for brain recordings, after which they were in neighbouring enclosures, separated by a metal grill as a precaution to prevent damage to their brain implants due to aggressive interactions. For a few days before the brain recordings, both monkeys were paired for about 30 minutes each day, to reconfirm their compatibility. On the day of recording, we paired them and allowed them to interact freely and naturally without any human intervention. They interacted intensively for ∼11 minutes while we recorded their brain activity simultaneously. After this, they stopped engaging with each other and we stopped the recording. Unfortunately it was only possible to record from one such social interaction session since the monkeys participated in many other solo recording sessions involving other behavioral tasks.

For video analysis, we selected one video camera feed in which most joints of both monkeys were clearly visible (netcam e3v817d, top view, 30 fps; Figure 5A). We used this video feed to manually mark the joint positions and used the CCTV camera videos to mark the times of all relevant social behaviors such as mounting, grooming and licking. We also marked the times at which each hand was used by each monkey. In addition, we manually annotated the joint positions of each monkey clearly visible in each frame throughout the session at 7-frame (0.233 s) intervals (joints annotated: head, neck, hip, right/left shoulder, elbows, wrists, palms, side-hips, knees, ankles and feet as well as the logger on the head; all annotated using NAPARI software). Joints that were not visible were marked as NaN (not a number). For all further analyses, we subtracted from the (x, y) position of each joint the (x, y) coordinates of the respective monkeys’ wireless logger, to obtain the joint position of each joint relative to the monkey’s own body.

The total duration of the social recording was 682 seconds. We observed unusually high firing in many PMv neurons of monkey M2 from 0-130 s of the session start. Since this unusual high firing could potentially confound the results, we removed this period from all further analyses. All results are shown with zero indicating start of analysis unless specified otherwise. We obtained qualitatively similar results even upon including this initial window. Similarly, we observed anomalous high-amplitude fluctuations in multi-unit activity (MUA) or local field potential (LFP) from some electrodes, and therefore excluded these from further analyses. All analyses in this study are based on 243 IT neurons (118 in M1, 125 in M2), 185 PMv neurons (57 in M1, 128 in M2) and 61 PFC neurons (all from M1). Likewise, all LFP analyses are reported from 215 IT sites (106 in M1, 109 in M2), 153 PMv sites (43 in M1, 110 in M2) and 60 PFC sites (all in M1).

### Touchscreen tasks

In order to interpret the neural activity during the social interaction as being of sensory or motor origin, we trained each monkey to perform a visual task and a motor task on a touchscreen attached to a juice spout, with custom-built eye tracking described previously (Jacob et al., 2021).

### Visual task

The visual task consisted of an eye calibration block and two fixation blocks. In the first eye calibration block, the monkey had to initiate the task by touching a circular button (4° radius) kept at (28°,0°) relative to the center of the screen within 10 s of its appearance. After this, four gray circles of 0.5° radius appeared sequentially (in random order) at four corners of a square 8° away from the center of the screen. Monkeys had to fixate each gray circle for 0.3 s successively, after which they received a juice reward. The trial was aborted if the monkey failed to fixate on the gray circle within a stipulated time (0.5 s). We recorded two correct trials of the calibration block. This data was used to learn a simple linear mapping between the eye tracker signal (x & y, arbitrary voltage units) and the touchscreen gaze location (x, y in degrees). We used this mapping to transform eye signal values in the fixation block to degrees of visual angle.

In the fixation blocks, the monkey had to start the trial by touching a blue button (4° radius) at (28°, 0°) relative to the center of the screen. Following this, a yellow fixation dot (0.1° radius) appeared at the center for 0.5 s. Upon successfully fixating for 0.3 s within a fixation window (10° radius), a sequence of 8 images was presented at the center of the screen, with each image appearing for 0.2 s followed by a blank screen of 0.2 s. If the monkey successfully maintained fixation within a certain window and maintained touch on the hold button within a certain window till the end of the trial, the trial was marked correct and the monkey was rewarded with juice (∼0.15-0.2 ml). Error trials were repeated after a random number of other trials. Since each monkey had different patterns of looking and touching, we used different hold button buffer windows (8° for M1, 2° for M2) and fixation windows (10° for face images for M1 and M2, 22° for body images for M1 and M2). Although the fixation window was kept relatively large, post-hoc analyses revealed that both monkeys eye positions were closely centered on the screen center throughout the trial (standard deviation of eye position during a trial: 0.65° for horizontal and 0.51° for vertical for M1, and 0.78° for horizontal and 0.70° for vertical for M2, averaged across trials; mean jitter in eye position across trials: 0.1° for horizontal and 0.22° for vertical for M1, and 0.2° for horizontal and 0.13° for vertical for M2).

The stimuli in the fixation block comprised photographs of 3 monkeys from the facility - M1 (*Co*), M2 (*Di*) & M3 (*Tu*) – that were photographed from different viewpoints. Monkey M3 was also group housed with the two monkeys before they were implanted. We selected five roughly matched views of each monkeys’ face (left, left-oblique, front, right and right-oblique), and five roughly matched views of each monkeys’ body in various poses (walking/sitting). Thus, there were 30 stimuli (3 monkeys x 5 face views + 3 monkeys x 5 body views). All the face images were scaled to measure 13.6° along the longest dimension, and body images were scaled to measure 43.8° along the longest dimension, to roughly match the size in real-world interactions. To avoid any potential neural response adaptation, successive images in a trial did not contain the same view or same monkey identity. Since image sizes were different for face and body images, they were presented in two separate blocks. Each monkey performed 60 correct trials of the fixation block, yielding 16 repetitions of each image which we used for subsequent analyses.

### Motor task

The motor task consisted of a calibration block which was the same as before, and a reach block. Each monkey performed the same task twice, once using his left hand and once with his right hand. We made monkeys switch their hand by fixing a metal grill on the left/right side of the juice spout that made it inconvenient to use the left/right hand to perform the task. Both monkeys were trained extensively to use the right hand for many other tasks, but adapted readily to use the left hand when the grill was placed to block access to their right hand.

In the reach block, each trial started with a green button that appeared 15° from the center on the left or right. After touching the button for 0.3 s, a second button appeared at one of four possible locations (up/down/left/right) equidistant (13°) from the center. The monkey had to touch the target button as soon as possible, and he received juice for correctly touching the button. Each monkey had to perform 64 correct trials (4 reach directions x 16 repeats). Error trials were repeated after a random number of other trials. For each monkey, we recorded separate sessions on the same day for the left and right motor tasks.

### Correlation with grooming surplus (Figure 2)

To compute the correlation between neural activity and the grooming surplus, for each time windows of 30 s each throughout the social interaction, we calculated the cumulative time spent by each monkey in grooming its partner minus the cumulative time spent receiving grooming. Likewise, we calculated the average firing rate of each neuron in these time windows. The correlation reported in Figure 2 is the Pearson’s correlation coefficient between the grooming surplus and the average firing rate across neurons (Figure 2C), or between grooming surplus and the firing rate of each neuron (Figure 2D).

### Grooming state decoding (Figure 3)

To investigate if population neural activity can be used to decode the current grooming state, we used a linear decoding analysis. We calculated the neural activity in successive 0.5 s bins across all neurons to form a vector of neural activations, and for each such bin, we assigned a 3-way label based on the corresponding grooming state in that bin (giving grooming, receiving grooming or no grooming). Bins containing a transition from one grooming state to another were excluded. To train the decoder, we first projected the multidimensional vectors corresponding to all bins into a low-dimensional space using principal component analysis by selecting principal components to capture 90% of the variance. We then trained the classifier using 5-fold cross-validation (*classify* function, MATLAB): in each training fold, we trained the classifier on 80% of the bins, and tested it on the held-out 20% of the bins. We concatenated the decoding accuracy for each held-out 20% of the data to obtain the cross-validated decoding label for all bins. By comparing this cross-validated decoding label with the actual label for each bin, we obtained a binary vector containing 0s for bins when the decoder was wrong and 1s for bins when the decoder was correct. Grooming decoding accuracy was calculated as the mean and standard error of this vector across bins.

### Partner identity decoding (Figure 4)

To decode the identity of the social partner, we first trained a 3-way linear decoder (using the same approach as the grooming state decoding) on the population neural activity from IT in the visual task elicited by the faces of M1, M2 and M3 during a 0-0.2 s window after image onset. The cross-validated decoding accuracy on the fixation task is shown using the red lines in Figure 4B-C.

To investigate whether this decoder, trained on the responses to face images on the touchscreen, would correctly decode the social partner’s identity during the social interaction, we calculated the population neural response vector during the social session in successive 0.5 s bins, centred each vector by subtracting the average firing rate of the corresponding grooming state and projected them into the principal components in the touchscreen task and obtained the predicted labels from the classifier. We excluded bins with transitions between the grooming bouts since those labels are ambiguous. The decoder was deemed correct in that bin and scored as 1 if the decoded identity was that of the social partner (i.e. for M1, social partner would be M2, and vice-versa) and scored as 0 if the predicted label was incorrect. Partner decoding accuracy was calculated as the mean and standard error of this vector across bins.

### Neural tracking of self & partner joint positions (Figure 5A-D)

To investigate whether single neurons were tracking the joint positions of the monkeys’ own joints (self) and that of its social partner, we first selected a subset of joints for further analysis. Of the 20 manually annotated joints, we found that upper body joints such as head, neck & shoulders (left & right) of both monkeys were visible in 79.3% of the frames, whereas other joints such as ankle and knee were frequently occluded or not clearly visible. We therefore selected these four joints (head, neck, left & right shoulder) for further analyses (Figure 5A). Including more joints into the model resulted in less data to fit the model, but yielded qualitatively similar results. Of the 2366 camera frames available from the manual annotations, we took 1876 camera frames with all these joints visible in both monkeys for further analyses.

For each neuron in each monkey, we calculated the firing rate in successive bins of the same width as the camera frame intervals centered on the camera frame times, and asked whether the firing rates across all these bins can be predicted as a weighted sum of the joint positions of itself, or its social partner. This amounted to solving a set of linear equations of the form ***y*** = ***Xb***, where ***y*** is an 1876 x 1 vector of firing rates of the neuron in each time bin across all three grooming states, ***X*** is a 1876 x 9 matrix containing the (x, y) coordinates of the head, neck, left and right shoulders and a constant, and ***b*** contains the weights of the linear summation. To evaluate this model, we used 10-fold cross-validation: we trained the model on 90% of the bins, and tested on the held-out 10% of the bins repeatedly to obtain a cross-validated prediction for all bins. Model performance was taken as the correlation coefficient between the predicted model output obtained in this manner with the observed firing rate, performed across bins of each grooming state (giving & receiving grooming). In this manner, we obtained the self joint model correlation (i.e. how well the neural response was predicted by the monkey’s own joints) and the partner joint model correlation (i.e. how well the neural response was predicted by the social partner’s joints) for each grooming state, which is shown in Figure 5B.

We note that these correlations are relatively low on average because the firing rates are based on discretizing the response into small time bins without the benefit of collecting multiple trials. We observed similar levels of correlation when we fit single trial firing rates in the visual task as a weighted sum of convolutional neural network (VGG- 16) activations, but we found much higher correlations upon fitting the same model to firing rates averaged across trials (data not shown).

### Neural mirror tuning for self & partner joints (Figure 5E-F)

To investigate whether neurons show similar tuning for self and partner joints, we created a combined model for each neuron in each monkey, where the neural response is taken as a weighted sum of the joint positions of its own joints as well as its partner’s joints (Figure 5E). The resulting model weights capture the tuning to both self joints (n = 8 weights) and partner joints (n = 8 weights) in the same model. To calculate model weights, we fit the model on firing rates across all bins, and calculated the correlation between the weights of the self joints and the partner’s joints. A positive correlation implies that a neuron that increases its firing for the monkey’s own joints also increases its firing when the partner’s joints move in the same way – in other words, this is the classic mirror neuron response. By contrast, a negative correlation means that the neuron that shows increases in firing for its own joints will decrease its firing when the partners joints make a similar movement – such activity would avoid direct production of movement upon observing one’s partner, yet enable understanding of the partner’s movements. These correlations are reported in Figure 5F-G.

### Interbrain coupling (Figure 6)

To investigate interbrain coupling, we asked whether the firing rate of each neuron in successive 0.05 s bins could be predicted as a weighted sum of the firing rates of neurons in the partner’s brain. In other words, we solved a linear regression of the form ***y*** = ***Xb***, where ***y*** is a 11,044 x 1 vector of firing rates of the neuron, ***X*** is a 11,044 x (n+1) matrix containing the firing rates of all *n* neurons in the partner monkey along with a constant term (set to 1), and ***b*** is a (n+1) x 1 weight vector that captures the contribution of each partner monkey neuron. To evaluate this model, we used a leave-one-out cross-validation: for each bin, we trained the model on all other bins and calculated the predicted firing rate in that bin. We built separate models for IT separately & PMv/PFC. Model performance for each neuron was evaluated as the correlation coefficient between the predicted firing rate obtained in this manner and the observed firing rate across all bins of the respective grooming state. To elucidate the time course of this partner model correlation, we computed the instantaneous correlation at each time bin across all neurons. Further, we smoothened the data by taking the average correlations in a 5 s moving window with a step size of 1s, so that we can capture the dynamics in a reasonable window. This model correlation (mean ± sem) is reported in Figure 6D)

## ACKNOWLEDGEMENTS

We thank Anindya Sinha for valuable insights and discussions on primate social interactions, Dr. Tim Blanche for technical assistance with wireless recordings, Dr. Sebastian Chandu for assistance with neurosurgeries, Dr. Sumedh Shastry for assistance with physiology during neurosurgeries and post-surgical care, Dr. Sumedh Shastry & Mr. V. Ramesh for primate facility management and animal care, Ravi & Ashok for animal care and Dharshana Raj & Vera Rao for assistance with manual annotation of joint positions.

## FUNDING

This research was supported by the DBT/Wellcome Trust India Alliance Senior Fellowship (Grant# IA/S/17/1/503081), Indian Institute of Science internal grants, a Google Asia Pacific research grant, and an intramural grant from Pratiksha Trusts Initiatives (all to SPA).

We also gratefully acknowledge the following fellowship support from the Government of India: Simon, S: Ministry of Education (MoE); Bhadra, D: Ministry of Education (MoE); Saha, S: Prime Ministers’ Research Fellowship (PMRF ID: 0200402) and Ministry of Education (MoE); Munda, S: CSIR Fellowship (ID: 35100868); Das, J: UGC Fellowship (816/CSIR-UGC NET).

## DATA AVAILABILITY

All data and code required to reproduce the results are publicly available at https://osf.io/tu5ed

## SUPPLEMENTARY MATERIAL

### SECTION S1. NEURAL RESPONSES DURING SOCIAL & TOUCHSCREEN TASKS

We have found that both IT & PMv neurons show elevated firing rates while giving compared to receiving grooming (Figure 3C). This could be due to increased sensory inputs and motor movements during giving compared to receiving grooming. To investigate this possibility, we compared neural responses in giving vs receiving grooming with the responses of these neurons during a visual task and a motor task performed on a touchscreen (see Methods; Figure S1A).

#### Responses of visual & non-visual neurons in IT while giving & receiving grooming

We hypothesized that, if there is greater sensory input while giving grooming, IT neurons that are visually responsive in a fixation task on the touchscreen should also show stronger modulation while giving vs receiving grooming. We therefore manually categorized each IT neuron in the visual task as visual or non-visual based on whether it showed a clear visual response to at least one image. We then calculated the average response of the visual and non-visual neurons during a response period (0.05-0.2 s after image onset) and a baseline period (−0.05 to 0.05 s relative to image onset). As expected, by definition, we observed a strong response modulation in the visual neurons compared to the non-visual neurons (Figure S1B). Next we compared the responses of these visual and non-visual neurons during the giving & receiving grooming bins of the social interaction. Interestingly, the neurons that show strong response modulation to visual images, also showed a strong response modulation to giving compared to receiving grooming (Figure S1F). Even the non-visual neurons in IT also showed a significantly larger response during giving compared to receiving grooming, although this difference was considerably smaller compared to the visual neurons (Figure S1F; average of absolute response difference, abs(giving – receiving grooming): 1.68 ± 0.16 spikes/s for visual neurons, 1.03 ± 0.22 spikes/s for non-visual neurons, p <0.05, ranksum test). These findings suggest that there is greater visual input during giving compared to receiving grooming. However, this analysis is not entirely conclusive since there are other signals in IT cortex during the social interaction that are not entirely sensory, as evidenced by the correlation with grooming surplus.

#### Responses of motor & non-motor neurons while giving & receive grooming

Next we hypothesized that, since there are greater motor movements during giving compared to receiving grooming, PMv neurons that are modulated by hand movements on touchscreen motor tasks should also show greater modulation for giving grooming. To this end, we manually categorized each PMv neuron in the motor task as being modulated by the movement and calculated their average response around a window relative to onset of response (0 to 0.3 s for the left hand, −0.5 to 0 s for right hand) compared to a baseline period (−0.35 to 0 s relative to sample onset).

This revealed a clear modulation in the motor task (Figure S1C-D), which is expected by definition. Upon comparing the responses of these two groups of neurons in the social session during giving and receiving grooming, we found that the motor neurons in PMv showed a clear response modulation (Figure S1G). This modulation was present strongly even in the non-motor neurons (Figure S1G), suggesting that the PMv neurons that were deemed non-motor on the touchscreen task based on only hand movements are presumably modulated by movements of other limbs in the social task. Nonetheless, the response difference in the motor neurons was larger than in the non-motor neurons (Figure S1G; average absolute response difference, abs(giving – receiving grooming): 4.35 ± 0.48 spikes/s for motor neurons, 0.99 ± 0.12 spikes/s for non-motor neurons, p < 0.00005, rank-sum test). These findings suggest that there is greater motor modulation during giving compared to receiving grooming. However, this analysis is not entirely conclusive since there are other signals in PMv cortex during the social interaction that are not entirely of motor origin, such as the correlation with grooming surplus.

We observed no clear pattern in PFC neurons – if anything, the non-motor neurons in PFC showed a larger response during receiving compared to giving grooming (Figure S1D, Figure S1H).

In sum, we conclude that the increased firing rates in IT & PMv during giving compared to receiving grooming are likely due to increased sensory inputs and motor outputs while grooming.

**Figure S1.**
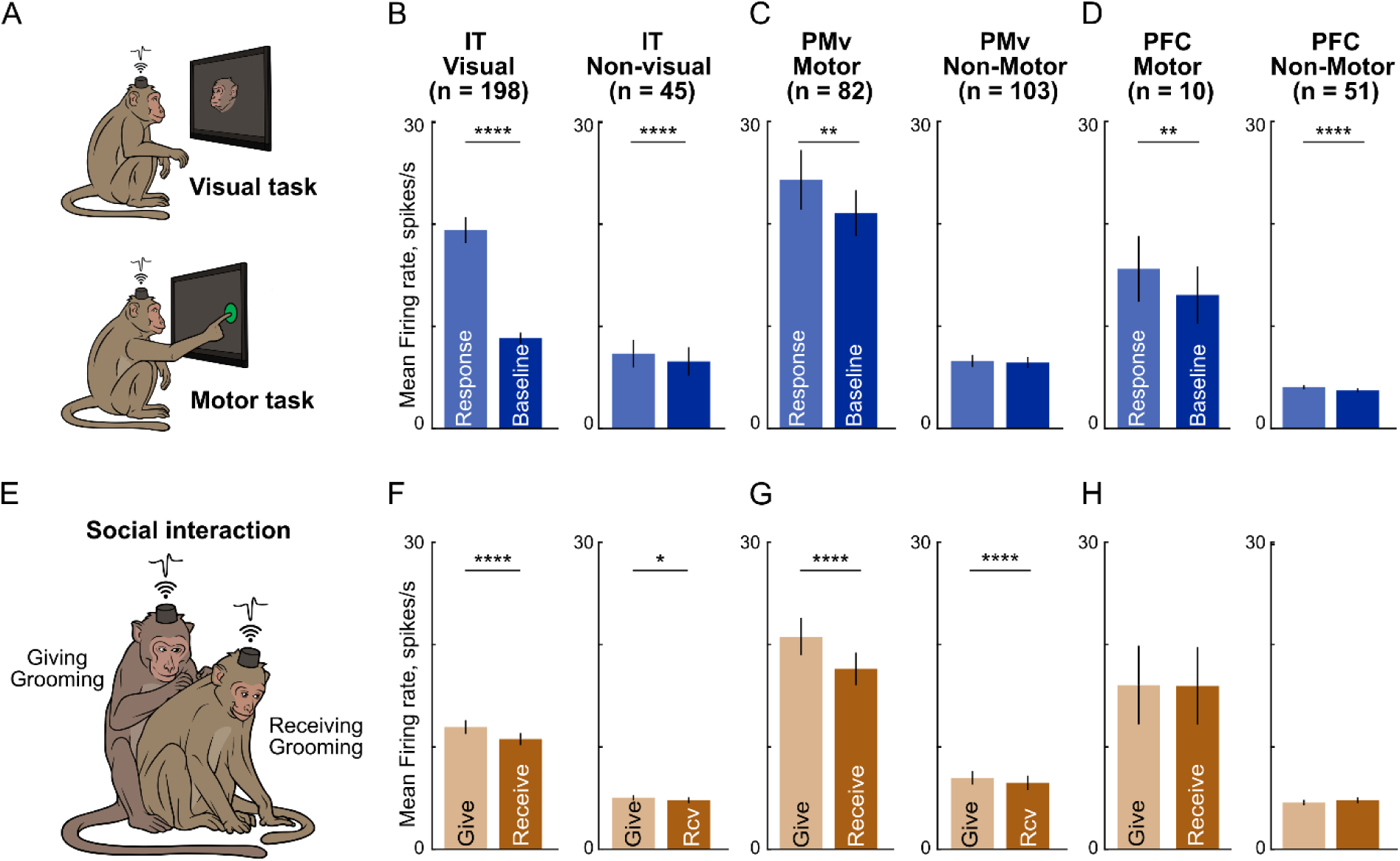
Visual & motor modulation during touchscreen and social tasks. (A) *Top:* Schematic of the visual task performed by each monkey separately on the touchscreen, where it had to fixate on a series of images presented visually. *Bottom:* Schematic of the motor task performed by each monkey separately on the touchscreen, where it had to touch targets appearing at four possible locations on the screen. (B) *Left:* Average firing rate of all visually responsive neurons in IT pooled across both monkeys during the image presentation period (0.05-0.2 s after image onset) and during the baseline period (−0.05 to 0.05 s relative to image onset). Asterisks represent statistical significance (* is p < 0.05, ** is p < 0.005, etc.; sign-rank test across neurons) *Right:* same plot for IT neurons classified as not visually responsive. (C) *Left:* Average firing rate of all motor-modulated neurons of PMv pooled across both monkeys aligned to the motor response and during the baseline period. *Right:* same plot for PMv neurons classified as non-motor. (D) Same as (C) but for motor & non-motor PFC neurons. (E) Schematic of social session, depicting M1 grooming M2. (F) Average firing rate for the visual (*left*) & non-visual (*right*) neurons in IT during giving and receiving grooming bouts. (G) Same as (F) but for motor & non-motor neurons in PMv. (H) Same as (G) but for motor & non-motor neurons of PFC.

### SECTION S2. ENCODING OF GROOMING SURPLUS VS OTHER SIGNALS

We have found that the average firing rate of IT neurons in each monkey closely tracks the grooming surplus signal (Figure 2). However, this correlation could arise from a simple overall level difference in firing rates between giving and receiving grooming. Likewise, monkeys might count the total number of giving versus receiving grooming bouts instead of keeping track of the overall duration. More generally, the correlation we have observed between neural responses and the grooming surplus signal could be alternatively explained by other factors that also vary in other ways during this period.

To this end, we created a number of variables that could covary with the average firing rate, as listed below. These signals are shown for each monkey in Figure S2A.

1. Grooming Surplus Time (GST), which is the main grooming surplus signal, defined as the total amount of time of giving grooming minus the total time of receiving grooming.
2. Cumulative Giving Grooming Time (CGGT), defined as the cumulative time spent in giving grooming to the partner.
3. Cumulative Receiving Grooming Time (CRGT), defined as the cumulative time spent in receiving grooming from the partner. Note that GST = CGGT – CRGT.
4. Cumulative Giving Grooming Number (CGGN), defined as the total number of bouts spent in giving grooming. Our rationale was that monkeys might keep track of the number of bouts instead of the cumulative time spent.
5. Cumulative Receiving Grooming Number (CRGN), defined as the total number of bouts spent in receiving grooming.
6. Cumulative Grooming Surplus Number (GSN), defined as the cumulative number of bouts spent giving grooming minus receiving grooming. Note that GSN = CGGN – CRGN.
7. Is Giving Grooming (IGG), which is a binary signal (0/1) indicating if the monkey is grooming the partner
8. Is Receiving Grooming (IRG), which is a binary signal (0/1) indicating if the monkey is receiving grooming from his partner
9. Within Bout Cumulative Giving Grooming Time (WBCGGT), which tracks the cumulative time spent in grooming the partner within each bout.
10. Within Bout Cumulative Receiving Grooming Time (WBCRGT), which tracks the cumulative time spent in receiving grooming within each bout.
11. Within Bout Cumulative Grooming Time (WBCGT), which tracks the elapsed time in a bout regardless of giving or receiving grooming.
12. α-power, which tracks the overall arousal and attentional state of the animal. To obtain the alpha power, we calculated the spectrogram of the local field potential signal on each channel (*pspectrum* function in MATLAB, in 1 s non-overlapping bins), took the power in the 8-12 Hz range, and averaged this alpha-band power across all channels in a given brain region. We obtained similar results on using channel-wise alpha power.

#### Correlation of each grooming signal with neural responses

We calculated each of these signals after smoothing the data with a 30 s boxcar window to capture the slowly varying nature of the signals (varying this choice yielded qualitatively similar results). For each neuron, we calculated the correlation between the firing rate and each variable across time, and averaged this across neurons to obtain an average correlation for a given brain region.

The resulting correlations are shown in Figure S2B for IT neurons. We find that grooming surplus time (GST) has the highest correlation, along with Is Giving Grooming (IGG; Figure S2B). Note that Is Receiving Grooming (IRG), being negatively correlated with IGG, has a negative correlation with the neural firing of IT neurons (Figure S2B). A number of other signals were also significantly correlated with the average neural response in IT. In IT and PMv, the firing rate was inversely correlated with the alpha power (Figure S2B).

We obtained a different pattern of encoding in PMv neurons: here, Is Giving Grooming (IGG) had the highest correlation compared to all other factors (Figure S2C). This is expected since PMv neurons show larger firing rates during giving grooming (Figure S1G), since grooming involves motor movements. Finally, PFC neurons showed relatively lower correlations (Figure S2D) compared to PMv and PFC, presumably because of the smaller number of cells sampled, that too only in one monkey.

#### Partial correlation of each grooming signal with neural responses

To confirm that the correlation of a given factor with the average firing rate is not explained by other factors, we performed a partial correlation analysis, where we calculated the correlation of the average firing rate of each neuron with a given factor after regressing out the possible contribution of all other factors. Since some signals are a combination of two other signals (e.g. GST = CGGT – CGRT), we included only two of these signals into the partial correlation analyses. In the resulting analyses, a significant partial correlation of a given factor implies that it has a unique correlation with the neural response even after considering all other grooming variables.

The resulting plots are shown in Figure S2E-G. For IT neurons, we found that grooming surplus time (GST) had the highest partial correlation that was significantly different from all other grooming variables (Figure S2E). However, a number of other factors had a significant partial correlation with IT neural responses as well.

For PMv neurons, we find that IGG had the largest partial correlation with neural responses (Figure S2F), with a few other variables continuing to have a significant but smaller contribution. Finally, PFC neurons showed no significant partial correlation with any variables – however this could be due to the smaller sample of PFC neurons, that too only in one monkey. There was no significant partial correlation between firing rates and mean alpha power in any of the brain regions (Figure S2F).

In sum, we conclude that the grooming surplus time is encoded strongly by IT neurons along with several other grooming-relevant variables, which could form the neural basis of grooming reciprocity.

**Figure S2.**
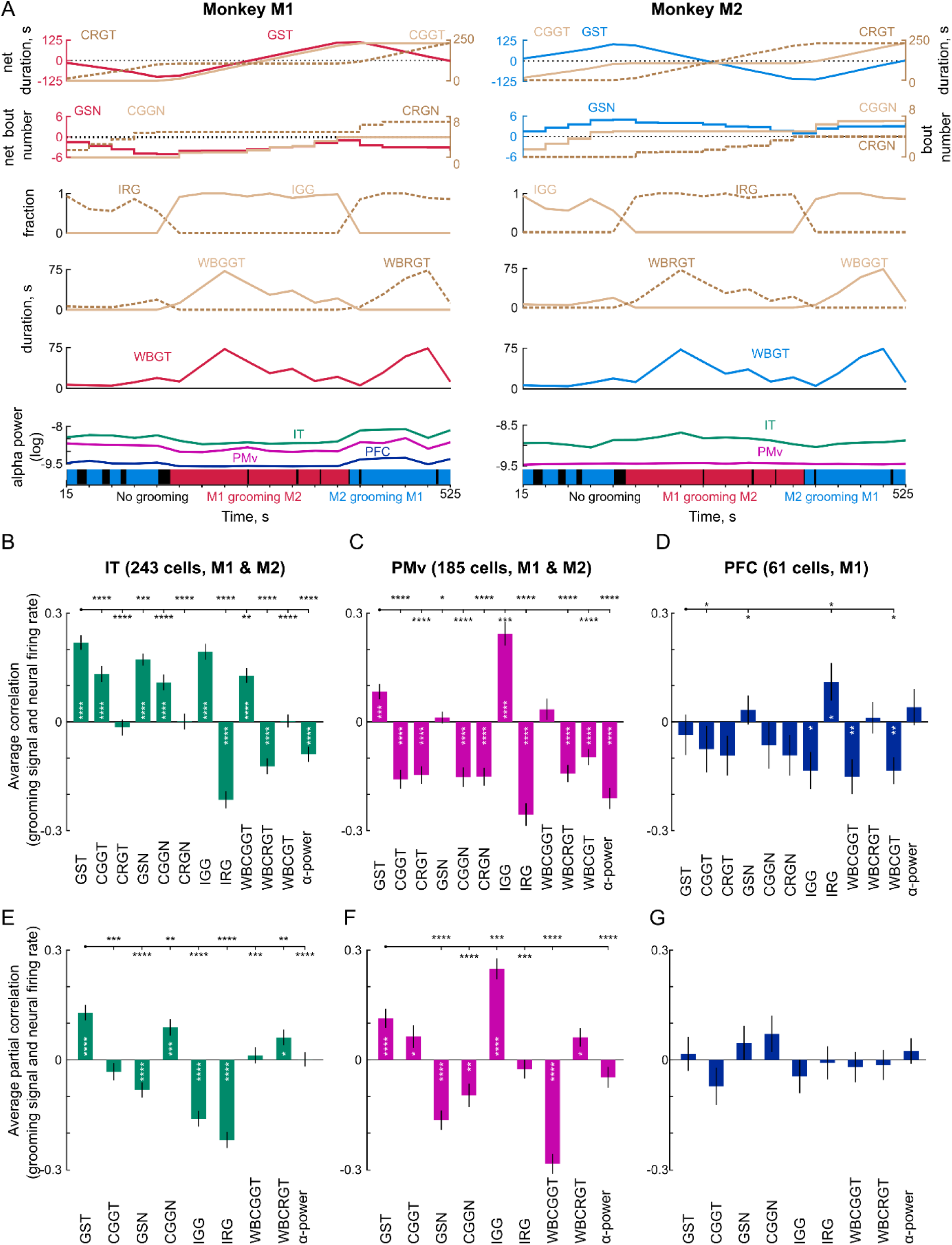
Encoding of grooming variables in each brain region. (A) Time course of grooming-related variables for monkey M1 (*left*) and M2 (*right*) that could be encoded by neural responses. Each grooming signal was smoothed using a 30 s window (we obtained qualitatively similar results on varying this choice). *First row:* Variables encoding cumulative time spent in grooming: Grooming Surplus Time (GST) overlaid with Cumulative Giving Grooming Time (CGGT) and Cumulative Receiving Grooming Time (CRGT). Note that GST = CGGT – CRGT. *Second row:* Variables that represent counting of bouts: Grooming Surplus Number (GSN) overlaid with Cumulative Giving Grooming Number (CGGN) and Cumulative Receiving Grooming Number (CRGN). *Third row:* Variables that represent binary state changes: Is Giving Grooming (IGG) and Is Receiving Grooming (IRG). *Fourth row:* Variables that represent elapsed duration only within each bout: Within Bout Cumulative Giving Grooming Time (WBCGGT) and Within Bout Cumulative Receiving Grooming Time (WBCRGT) *Fifth row:* Variables that represent elapsed duration of interaction regardless of give or receiving grooming: Within Bout Grooming Time (WBCGT). *Sixth row:* α-power (8-12 Hz), averaged across sites in IT (*green*), PMv (*magenta*) and PFC (blue), which is a proxy for the animals’ arousal/attentional state. (B) Average correlation (mean ± s.e.m across neurons) between each grooming variable with average firing rate of IT neurons (n = 243 from M1 & M2). White asterisks indicate statistically significant correlations (* is p < 0.05, ** is p < 0.005, etc., sign-rank test across neurons). Asterisks above each pair of bars indicate statistically significant comparisons (* is p < 0.05, ** is p < 0.005, etc., sign-rank test across neurons). (C) Same as (B) but for PMv (n = 185 across M1 & M2) (D) Same as (B) but for PFC of M1 (n = 61 in M1) (E) Average partial correlation (mean ± s.e.m across neurons) between each grooming variable and average firing rate after controlling for all other factors for IT neurons (n = 243 from M1 & M2). Asterisks follow the same convention as before. (F) Same as (E) but for PMv (n = 185 across M1 & M2) (G) Same as (E) but for PFC of M1 (n = 61 in M1)

### SECTION S3. SPIKE & LFP POWER SPECTRA ACROSS GROOMING TYPE

#### Introduction

We found that IT & PMv show elevated firing while each monkey is grooming his partner compared to when it receives grooming. Here, we investigated whether these firing rate increases are driven by increased power in specific frequency bands, by calculating the power spectrum as a function of frequency. In addition, we also investigated whether the increased firing rates during grooming are accompanied by changes in the frequency of local field potential oscillations in each brain region.

#### Methods

Local field potentials (LFP) were extracted from the wideband signal recorded from each electrode by band-pass filtering the signal in the window [0.3 500] Hz and down- sampling to 2500 samples/s., whereas the multiunit activity (MUA) was extracted from the wideband signal by using Plexon Offline Sorter. We observed high amplitude fluctuations in some of the electrodes in MUA & LFP data, and therefore removed them from further analysis. In all, we analyzed LFP signals from 215 electrodes in IT, 153 in PMv and 60 in PFC, and analyzed MUA from 243 neurons in IT, 185 in PMv and 61 in PFC across two monkeys.

We computed the power in the MUA and LFP signal from each electrode using the MATLAB function *‘pspectrum’* (using 1-second time resolution, no overlap), and averaged the power across all time bins of each grooming type (giving & receiving grooming) to obtain a global average power at each frequency for each electrode. MUA firing rates were obtained by calculating the spike counts in bins of 1/2500 s, so as to match the sampling rate of the LFP signal and make the spectra more comparable.

#### Results

The average MUA & LFP power spectra across all neurons for each brain region and each grooming type is shown in Figure S3. Below we describe the effects observed in each brain region.

#### Average MUA & LFP power spectra in IT

In monkey M1 MUA, we observed a decrease in power around the alpha band (8- 12 Hz) in giving compared to receiving grooming, and an increase in power at high frequencies greater than 24 Hz (Figure S3A). In monkey M1 LFP, we observed the opposite pattern (Figure S3B): there was decreased power in all frequencies during giving grooming, with the effect particularly pronounced in the alpha band and in around 39 Hz. Thus, there is increased alpha power during receiving grooming in both firing rate and local field potentials, which might be due to monkey M1 closing his eyes or relaxing while receiving grooming from his partner.

In monkey M2, we observed a wideband increase in MUA power in the giving compared to receiving grooming condition. This was accompanied by a selective decrease in alpha power in LFP and an increase in power around 63 Hz (Figure S3B). Thus, there is increased alpha power during receiving grooming in M2 but only in LFP but not in the firing rate.

Thus, the increased firing rate in IT during giving grooming is accompanied by decreased LFP power particularly in the alpha band. We speculate that this could be due to increased inhibitory synaptic input during receiving grooming, driven by alpha band oscillations.

#### Average MUA & LFP power spectra in PMv

In monkey M1 MUA, we observed consistently larger power in giving grooming at all frequencies (Figure S3C), with the smallest effect in the alpha band. By contrast, the LFP power was consistently smaller in giving grooming compared to receiving grooming, with the largest effect around 20 Hz (Figure S3D).

In monkey M2 MUA, there was uniformly larger power in giving grooming at all frequencies (Figure S3C). By contrast, the LFP power was smaller in giving grooming around 24 Hz and larger in higher frequencies around 42 Hz (Figure S3D).

Thus, the increased firing rate in PMv during giving grooming is accompanied by reduced LFP power particularly in the alpha band. We speculate that this could be due to increased inhibitory synaptic input during receiving grooming, driven by alpha band oscillations.

#### Average MUA & LFP power spectra in PFC

In monkey M1 MUA, we observed lower power during giving grooming at all frequencies (Figure S3E). This was accompanied by reduced LFP power during giving grooming compared to receiving grooming, with the effect most prominent in the alpha band (Figure S3F). We speculate that this could be due to increased PFC activity during receiving grooming when the monkey is directing the grooming of his partner towards desirable locations.

#### Dynamics of MUA & LFP spectra across brain regions

To characterize the dynamics of these spectral power variations across time, we visualized the power spectrum for each bin separately for MUA (Figure S4) and for LFP signals (Figure S5). It can be seen that there are specific times at which the spectral power is modulated even within grooming bouts, and that there is greater spectral power during giving grooming in MUA (Figure S4) and greater spectral power during receiving grooming in LFP (Figure S5).

#### Conclusions

We conclude that both IT & PMv show increased firing rates during giving grooming, which is accompanied by decreased LFP power, particularly in the alpha band. We speculate that this could be due to increased inhibitory synaptic input during receiving grooming, driven by alpha oscillations.

**Figure S3.**
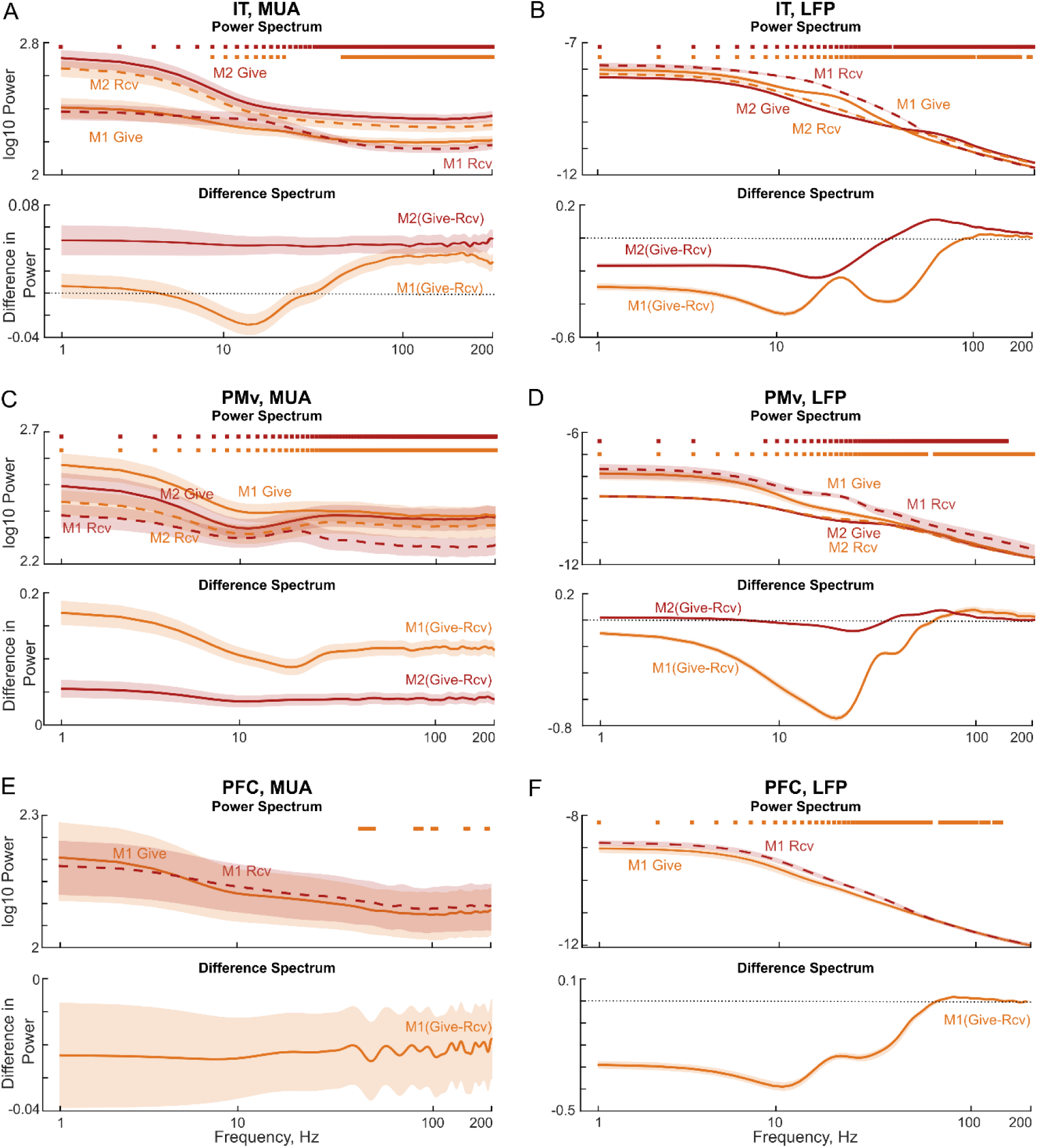
Multiunit activity & local field power spectra across grooming types. (A) *Top:* Average power spectrum of multiunit activity (MUA) across all IT neurons in give & receiving grooming conditions in M1 (n = 118) and M2 (n = 125). Shaded regions indicate s.e.m across neurons. The squares at the top indicate statistically significant differences in power at each frequency (sign-rank test, p < 0.05). *Bottom:* Difference in the power between give and receiving grooming for M1 & M2 (B) Same as A, but for local field potentials (LFP) in IT cortex from M1 (n = 106) and M2 (n = 109) (C) Same as A, but for MUA in PMv in M1 (n = 57) and M2 (n = 128). (D) Same as B, but for LFP in PMv cortex of M1 (n = 43) and M2 (n = 110) (E) Same as A, but for MUA in PFC of M1 (n = 61). (F) Same as B, but for LFP in PFC of M1 (n = 60).

**Figure S4.**
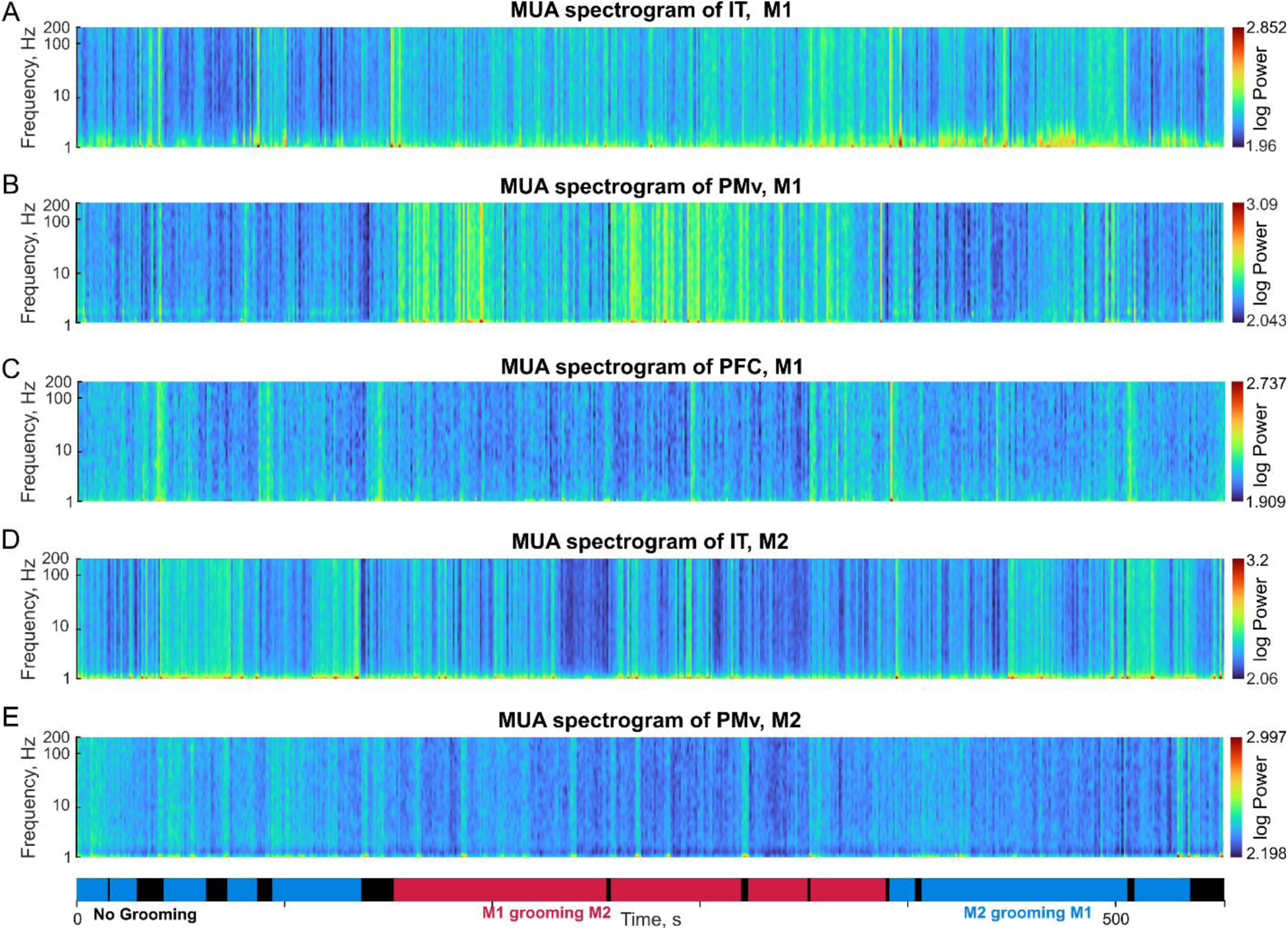
MUA spectrogram during the social interaction. (A) Spectrogram of the multiunit (MUA) activity in IT cortex of monkey M1 averaged across neurons (n = 118). Frequency is shown along the y-axis on a logarithmic scale. The plot on the bottom shows the timeline of the social interaction with the three types of grooming behaviors: M1 grooming M2 (*red*), M2 grooming M1 (*blue*) and no grooming (*black*). (B) Same as (A) but for PMv of M1 (n = 57) (C) Same as (A) but for PFC of M1 (n = 61) (D) Same as (A) but for IT of M2 (n = 125) (E) Same as (A) but for PMv of M1 (n = 128)

**Figure S5.**
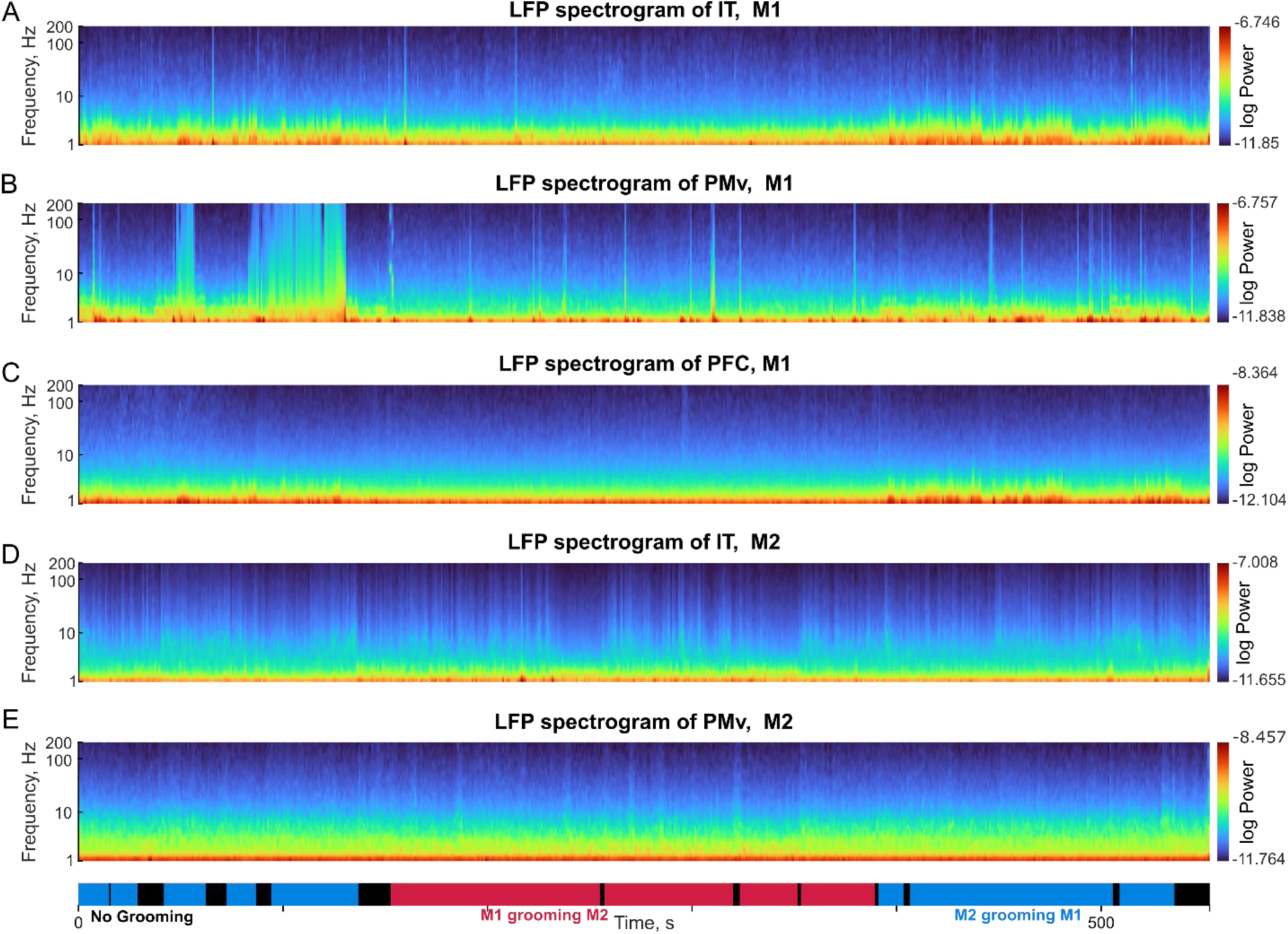
LFP spectrogram during the social interaction. (A) Spectrogram of local field potential (LFP) activity in IT cortex of monkey M1 averaged across neurons (n = 106). Frequency is shown along the y-axis on a logarithmic scale. The plot on the bottom shows the timeline of the social interaction with the three types of grooming behaviors: M1 grooming M2 (*red*), M2 grooming M1 (*blue*) and no grooming (*black*). (B) Same as (A) but for PMv of M1 (n = 43) (C) Same as (A) but for PFC of M1 (n = 60) (D) Same as (A) but for IT cortex of M2 (n = 109) (E) Same as (A) but for PMv of M2 (n = 110)

### SECTION S4. GROOMING DECODING ACROSS TRANSITIONS

We found that neural activity can be used to decode the ongoing grooming state during the social interaction (Figure 3). While this is expected since each state involves distinct sensory inputs and motor outputs, a more interesting possibility is that neural activity also carries information about an upcoming transition from one grooming state to another.

To investigate this issue, we calculated the grooming decoder probabilities in successive 0.5 s bins in a 31 s window around each transition: from No Grooming to Giving grooming and vice versa (Figure S6A) and from No Grooming to Receive Grooming and vice-versa (Figure S6B). In all cases, we found a gradual transition whereby the decoder probability increased a few seconds before the transition itself. These changes could well be due to changes in behavior of the animals as they prepare for the transition, or due to preparatory neural activity that is commonly found to precede motor movements.

Thus, neural activity carries information about the future grooming state around the time of a transition.

**Figure S6.**
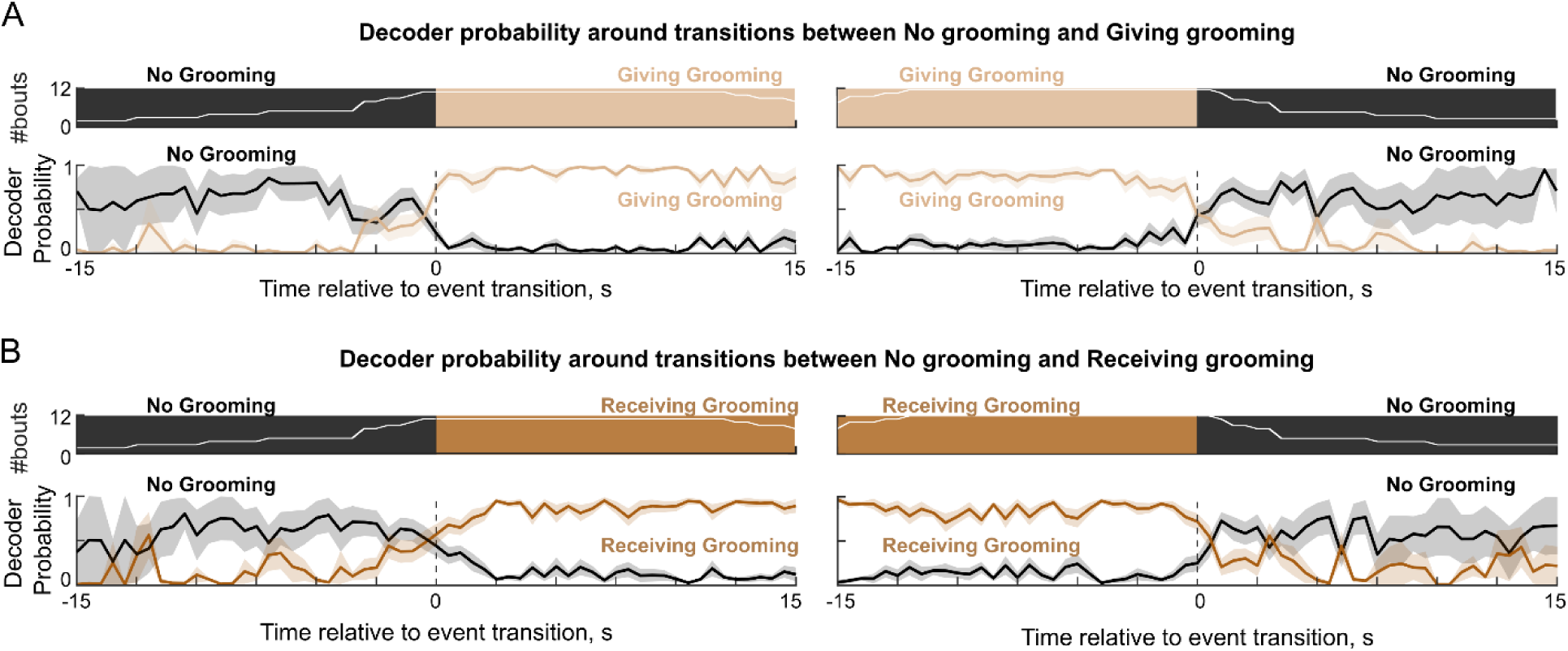
Grooming state decoding during transitions between grooming bouts. Average grooming state decoding probability around the time of transitions from one grooming type to another. The *top row* in each panel shows the number of bouts included at each point in time relative to the event transition, and the bottom row depicts the grooming decoder probability averaged across these number of bouts. Shaded bars represent s.e.m across bouts. (A) *Left:* Grooming state decoding probability for giving grooming (in brown color) and no grooming (in black color) around the transition from no grooming to giving grooming. *Right:* Same as earlier, but for the transition from giving grooming to no grooming. (B) Same as (A) but in place of giving grooming, receiving grooming probability is plotted for transitions from no grooming to receiving grooming (*left*) and from receiving grooming to no grooming (*right*)

### SECTION S5. DYNAMICS OF PARTNER AND HAND DECODING

#### Dynamics of partner identity decoding during social interaction

We have found that partner identity could be reliably decoded from IT neurons with no systematic differences across different type of grooming bouts (Figure 4). To investigate whether this decoding was stable throughout time, we calculated the face decoding accuracy as a function of time from both monkey M1 IT (Figure S7A) and monkey M2 IT (Figure S7B). We found that face decoding accuracy changed dynamically throughout the session, presumably higher when the monkeys were tracking each other.

#### Partner identity decoding during head vs body grooming

The above variations in face decoding could be due to the animals’ own social status, or due to the animals tracking each other more or less closely during the social interactions. To investigate this issue further, we wondered whether animals might be tracking their partners’ identity more or less closely when they are grooming their partners’ head compared to the body, or even when receiving grooming on the head compared to the body.

The results are shown in Figure S8. We observed interesting variations in partner decoding in monkey M1: here, partner decoding was slightly higher when M1 was grooming M2 on the head compared to the body (Figure S8A). Similarly, partner decoding was higher in M1 when M2 was grooming his head. Thus, M1 encoded his social partner’s identity strongly when grooming was either given or received on the head.

We observed a different pattern of partner identity decoding in M2: here, partner decoding accuracy in M2 was high regardless of whether M1 was giving grooming on the head or body, and even when M2 was grooming the head or body of M1 (Figure S8B). Thus, M2 strongly encoded his social partner’s identity regardless of head or body grooming.

#### Dynamics of hand decoding during the social interaction

Since each monkey performed a motor task with his left and right hand (see Methods), we wondered whether it would be possible to decode the hand that is being used during the social interaction. To this end, we trained a 2-way decoder on the motor task responses in each brain region using the same methods as before, and tested it during times during the social interaction when we annotated each monkey using his left or right hand (Figure S9A). We found that hand decoding was accurate only in PMv (Figure S9B).

To investigate the dynamics of hand decoding, we calculated the hand decoding probability across time using the PMv activity in each monkey. Since, in our manual annotation, we specified the left/right hand usage only when the monkey initiated the grooming, the hand labels are not available for the entire recording. Hence, we show the probability of left hand label, instead of accuracy (Figure S9 C,D). We smoothened the probability of left hand label, in a window of 5 s with a step size of 1 s. The mean and s.e.m. of the probability values in each 5 s window is shown (Figure S9 C,D top plots) This revealed dynamic variations in hand decoding probability which corresponded nicely to the left or right hand being used by each monkey for grooming during the social interaction (Figure S9 C-D).

Thus, there is a systematic relationship between neural activity evoked during more controlled motor tasks and the motor movements that occur during social interaction.

**Figure S7.**
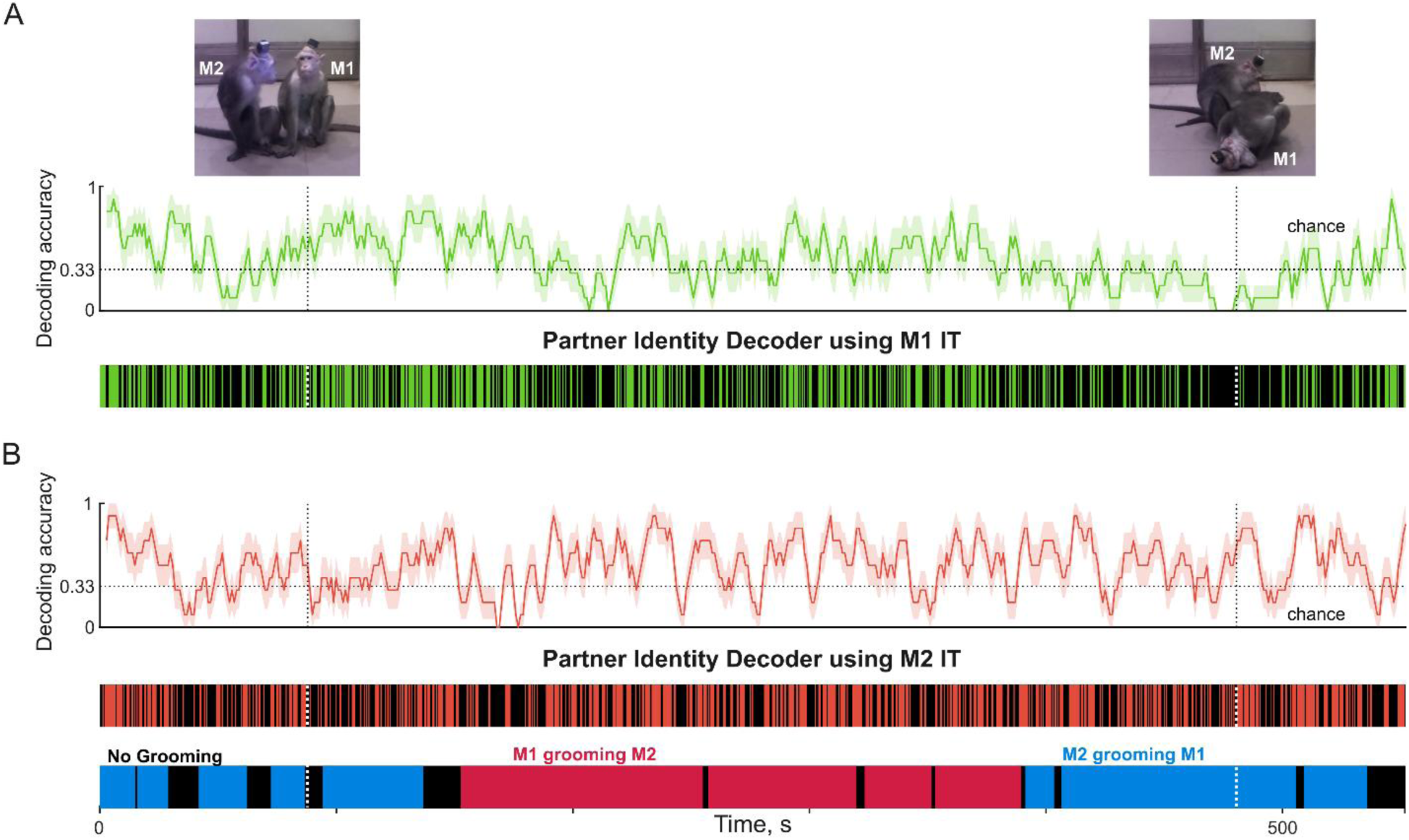
Dynamics of partner identity decoding during social interaction. (A) *Top:* Decoding accuracy of partner identity decoding, calculated across 5 s bins with a step size of 1 s from M1 IT neurons, with example frames illustrating the two monkeys when they are facing each other, and when they are not facing each other. *Bottom:* Decoding accuracy in individual time bins throughout the social interaction session (*green*: correct, *black*: incorrect) (B) Same as (A) but using monkey M2 IT neurons (*red*: correct, *black*: incorrect).

**Figure S8.**
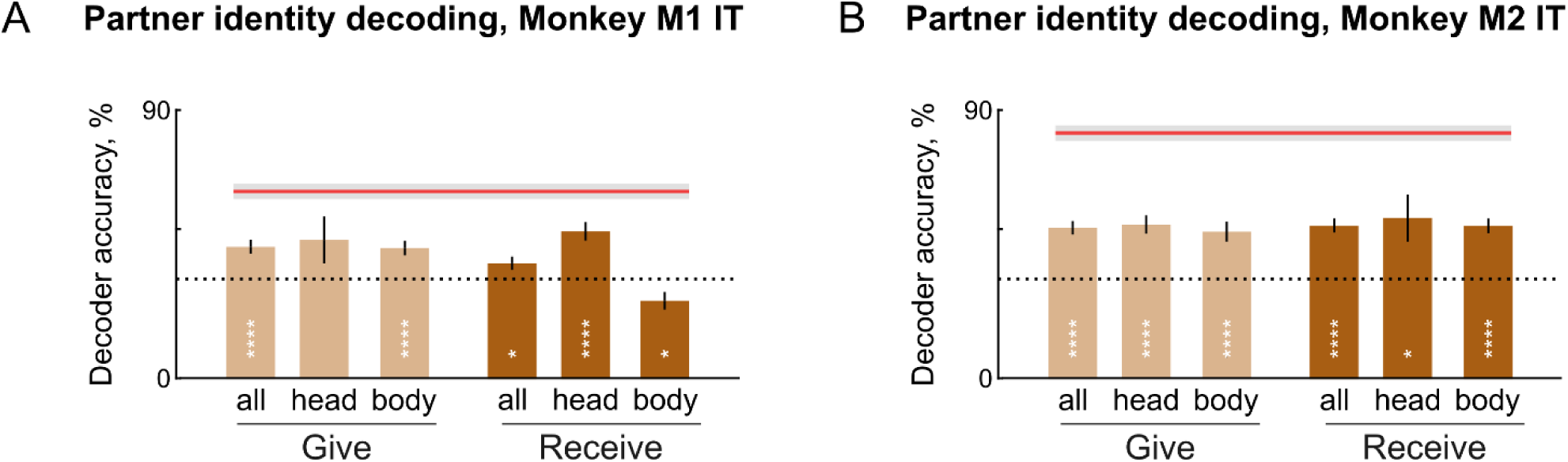
Partner identity decoding during head and body grooming. (A) Partner identity decoding accuracy from M1 IT neurons during giving (*left, light brown*) and receiving (*right, dark brown*) groom states, when the grooming was directed towards the head or the body. Decoding accuracy was calculated on non-overlapping 0.5 s bins, and each bin was uniquely assigned to either giving/receiving groom and head/body grooming. Error bars represent s.e.m. across all bins of that type. Decoding accuracy in the touchscreen task is also shown for reference (*red lines, with shaded regions* indicating s.e.m.). Asterisks represent statistically significant decoding accuracy (* is p <0.05, **** is p < 0.00005, chi-square test of correct and incorrect counts across bins compared to chance). (B) Same as (A) but using monkey M2 IT neurons.

**Figure S9.**
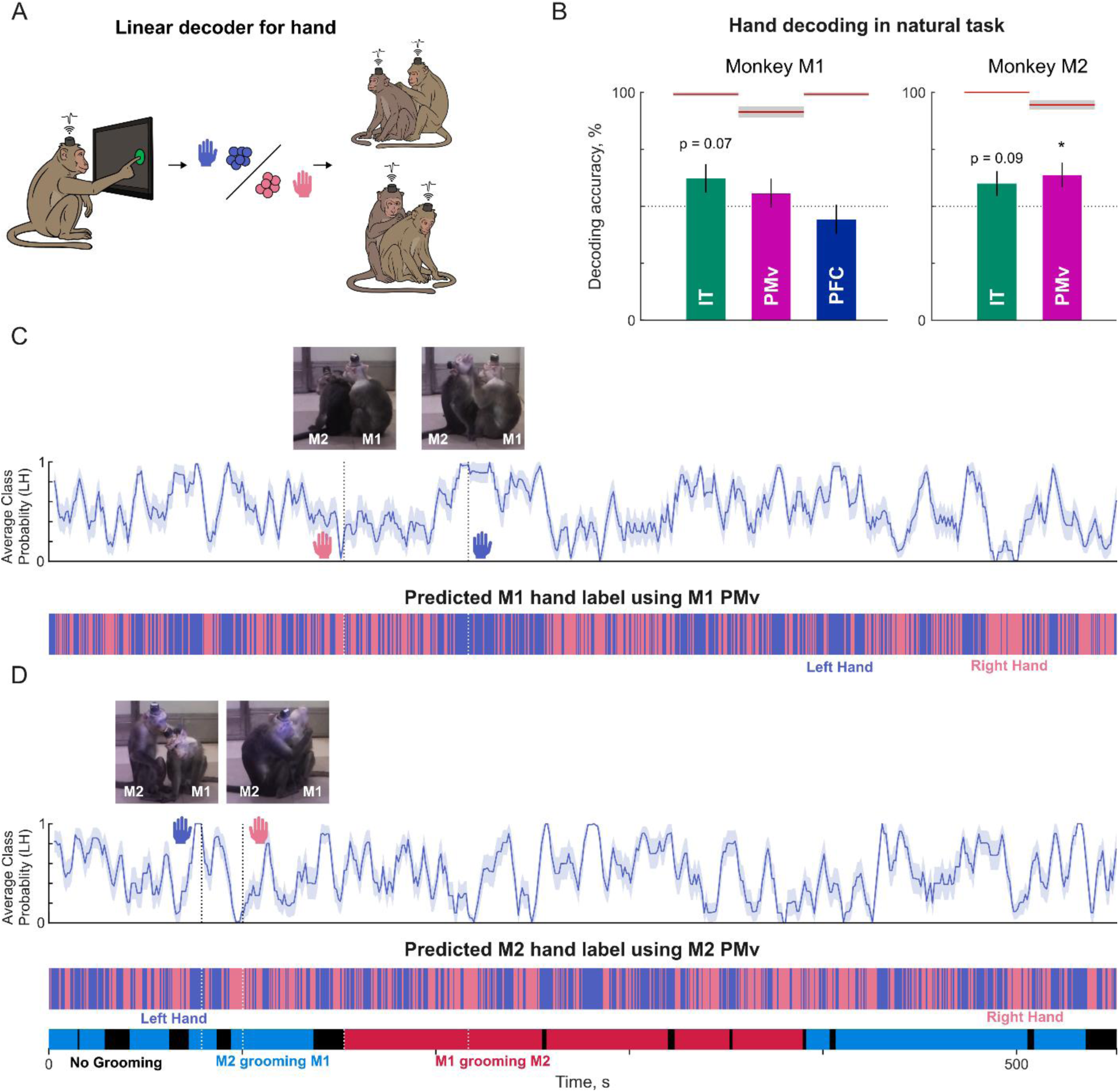
Dynamics of hand decoding during social interaction. (A) *Left:* Schematic of the motor task performed by each monkey separately, where he had to touch a target appearing on the screen with his left hand or right hand. *Middle:* We trained a two-way linear decoder on the neural responses on the motor task to decode which hand is being used by the monkey. *Right:* We tested the decoder during the social interaction. (B) Hand decoding accuracy from monkey M1 (IT, PMv & PFC) and from monkey M2 (IT, PMv). Red lines indicate decoding accuracy from the motor task, with shaded regions representing s.e.m across trials. Asterisks represent statistical significance of the decoding accuracy being above chance level (p < 0.05, chi-square test across 21 bins & 1 instances of left hand & 40 bins and 2 instances of right hand in M1 and 40 bins & 2 instances of left hand & 40 bins and 2 instances of right hand) (C) *Top:* Average class probability given by the decoder for ‘Left Hand’ label, calculated across 5 s bins with a step size of 1 s from M1 PMv neurons, with example frames illustrating various types of hand use. *Bottom:* Predicted hand label in individual time bins throughout the social interaction (*pink*: right hand, *blue*: left hand) **(D)** Same as (C) but using monkey M2 PMv neurons.

### SECTION S6. ADDITIONAL ANALYSIS OF SELF & PARTNER JOINT MODELS

#### Self and partner joint model correlations for different levels of joint motion

We have found that neural activity in each monkey is predicted better by the partner’s joint positions during giving grooming and by his own joints during receiving grooming (Figure 5). However, these self and partner joint correlations could be modulated by the amount of motion in these joints, which could be different between give and receiving grooming.

To investigate this issue further, for each joint, we calculated the motion energy by taking the absolute difference between the joint position of each bin with the next bin. We then calculated the z-scored motion signal from each joint position, and averaged it across joints to obtain an average motion signal. We then calculated the difference in cross-validated model correlation between self and partner joint models for different levels of self and partner motion (in quartiles). The resulting plots are shown in Figure S10.

We found qualitatively similar results for all levels of joint motion (Figure S10). In other words, during giving grooming, neural activity of the groomer was predicted better by the partner’s joints and vice-versa during receiving grooming. However, when the monkeys’ own joints had higher motion strength, we found that the self joint model predicted neural activity better (Figure S10).

#### Dynamics of self & partner joint correlations with time

To investigate the dynamics of the self and partner joint models across time for all neurons, we calculated instantaneous correlations between the observed and predicted firing rates of all neurons (after excluding the noisy ones). We averaged the instantaneous correlation values in 5 s time windows in time steps of 1 s. The time course of the difference between self and partner joint model correlations is shown for each region in Figure S11. We find distinct periods within each grooming bout in which the partner model has a higher correlation compared to the self-joint model, complementing our observation that the partner drives grooming in social interactions.

**Figure S10.**
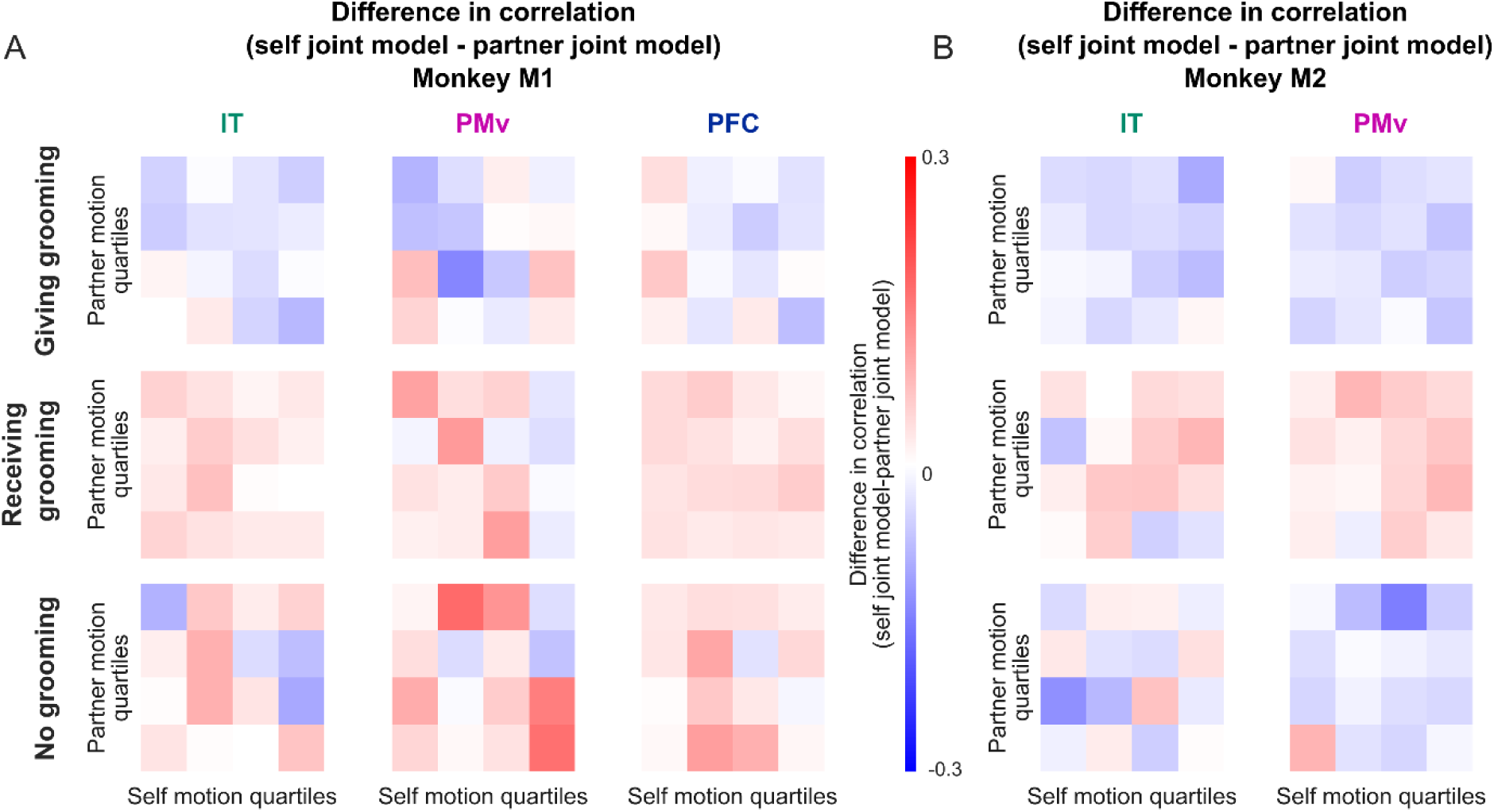
Self & partner model correlations for different levels of joint motion. (A) Difference between self and partner joint model correlations in IT (*left*), PMv (*middle*) and PFC (*right*) monkey M1 for giving grooming (*top*), receiving grooming (*middle*) and no grooming (*bottom*) bins of the social interaction. Each color map contains the difference in the model correlations for each quartile of self and partner motion, with *red* indicating self joint model correlation being larger than the partner model, and *blue* indicating the opposite. (B) Same as (A) but for monkey M2.

**Figure S11.**
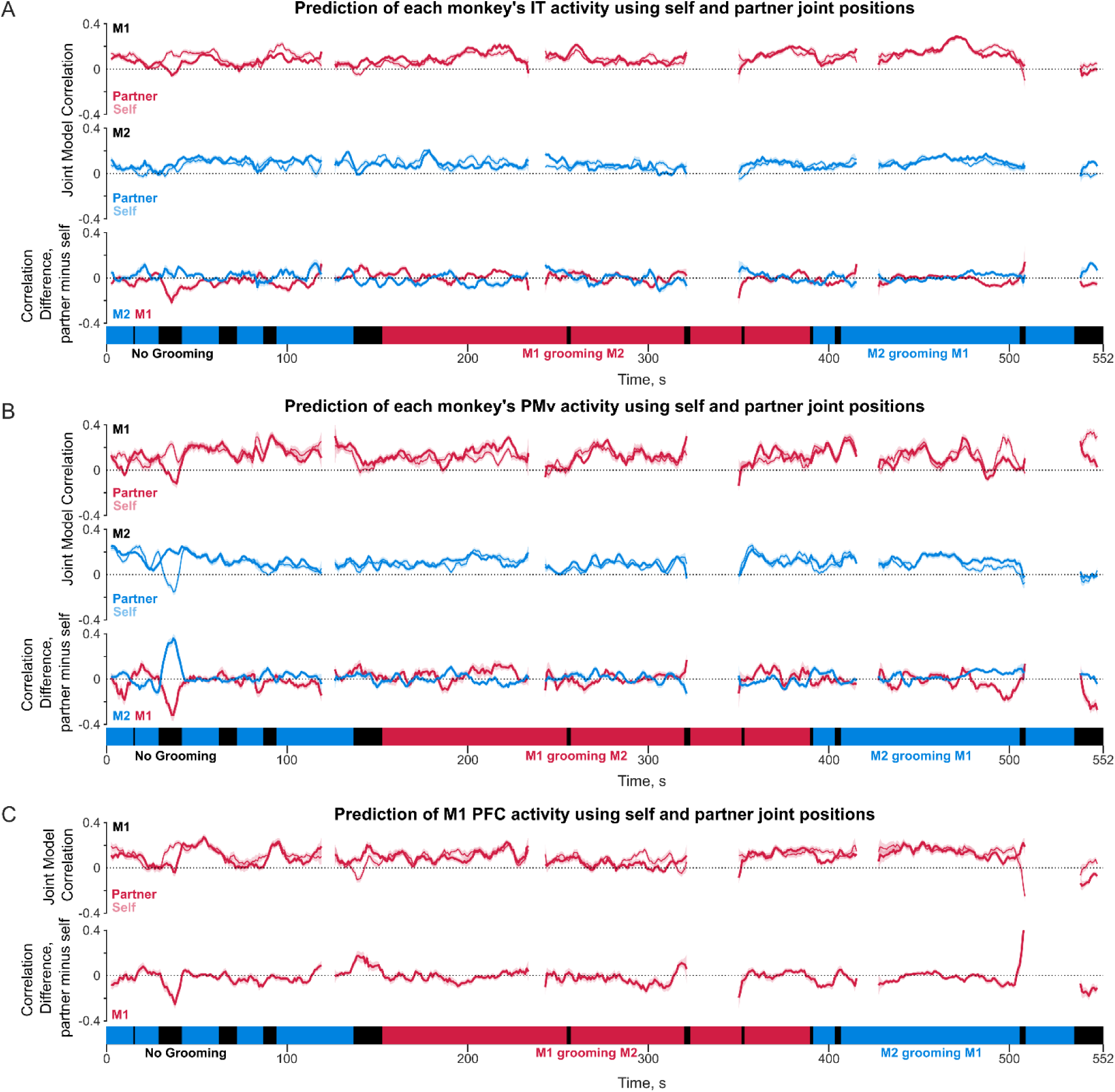
Time course of self & partner joint correlations during social interaction. (A) Top. Time course of smoothened instantaneous correlation of the partner *(darker)* and self *(lighter)* joint models in IT for M1, with shaded bars representing s.e.m across average correlation within moving window of 5s. Middle. Same as Top, but for M2. Bottom. Difference between partner and self joint model correlation plotted over time for M1 (red) and M2 (blue) separately. (B) Same as (A) but for PMv (C) Same as (A) but for PFC.

### SECTION S7. INTERBRAIN COUPLING DYNAMICS ACROSS BRAIN REGIONS

We have found that brain activity of the groomer is driven the receivers brain activity for PMv & PFC considered together. Here we report the interbrain coupling for each brain region separately (Figure S12).

In IT, we found that the brain activity of the groomer is predicted by the receiver only slightly better than the other way around, and this effect did not attain statistical significance (Figure S12A). Upon computing these partner model correlations as a function of time, we found distinct periods where the neural activity of each monkey is strongly predicted by his partner (Figure S12B). We speculate that these could be periods where both monkeys are looking at common visual items in their environment, or at each others movements.

In PMv, we found that the brain activity of the groomer is predicted by the receiver, and this effect was statistically significant (Figure S12C). Here too, we found the neural activity of each monkey to be strongly predicted at some phases during the social interaction compared to others (Figure S12D). We speculate that these could be periods when the monkey was closely tracking the movements of his partner.

Finally, in PFC, we have brain recordings only from M1, but here we find that PFC neurons were predicted better by M2 neural activity during M2 grooming M1 compared to when M1 groomed M2, but this did not reach significance (p = 0.93) Here too, we found neural activity of M1 to be strongly driven by his partner at some times during the social interaction compared to others (Figure S12F). We speculate that these could be periods of shared visual attention or goals, or shared tracking of each other’s movements.

Taken together, we find that interbrain coupling exhibits rich dynamics across the course of the social interaction, and these dynamics are largely driven by the receiver of the grooming.

**Figure S12.**
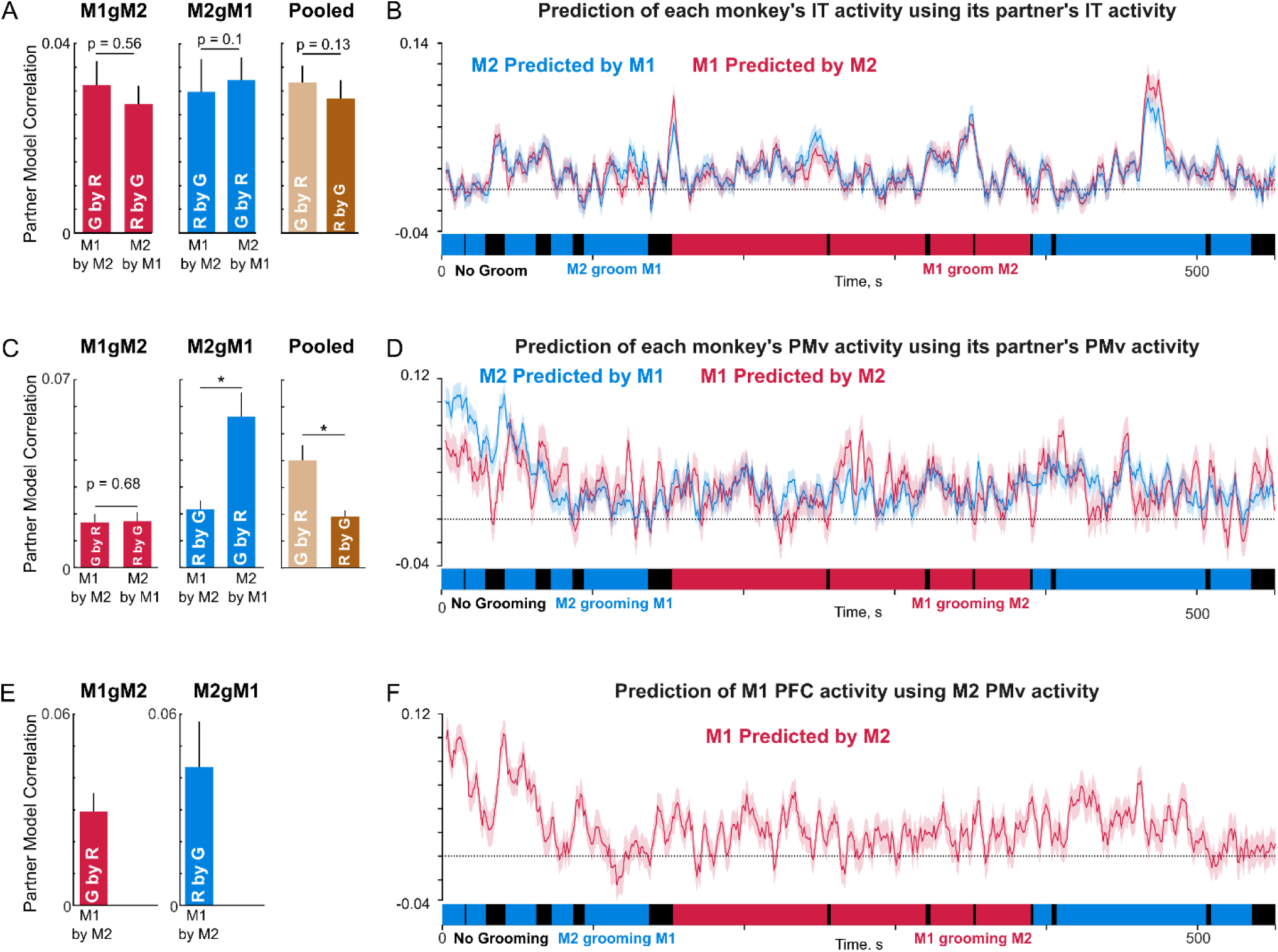
Dynamics of interbrain coupling across brain regions. (A) Partner model correlation (mean + s.e.m) in IT cortex of M1 & M2 while M1 groomed M2 (*left, red bars*), while M2 groomed M1 (*middle, blue bars*), and pooled data for the groomer and receiver monkey (*right, brown bars*), across the neurons which show significant model correlation. Asterisks indicate statistically significant comparisons (* is p < 0.05, Wilcoxon rank sum test across neurons). ‘G by R’ indicates the average correlation across neurons when groomer’s brain activity is predicted by receiver’s brain activity and ‘R by G’ indicates the average correlation across neurons when the receiver’s brain activity is predicted by the groomer’s activity. (B) Time course of partner model correlation during the entire interaction (mean ± s.e.m, solid lines with shaded regions across all 50 ms time bins in the 5 s smoothening window). (C-D) Same as (A) & (B) but for PMv cortex of M1 & M2. (E-F) Same as (A) & (B) but for PFC of M1.

### SECTION S8. INTERBRAIN CROSS-CORRELATION

We have found that the groomer’s brain activity is predicted by the receiver’s brain activity in both monkeys (Figure 6). However, this analysis is based on instantaneous activity and does not account for possible temporal relationships (like a lead or lag) between the two brains, or possible coherence in specific frequency bands. Here, we investigate these possibilities. We did not find clear patterns, but nonetheless included detailed analyses here for the sake of completeness.

#### Inter-brain cross-correlation dynamics during social interaction

We began by asking whether there is any temporal relationship between the neural activity in the two monkeys during the social interaction. For instance, the neural activity of the groomer could precede the neural activity of the receiver since the groomer initiates motor movements whose sensory consequences are registered in the receiver’s brain.

To this end, we computed the cross-correlation between every possible pair of neurons in the IT cortex of the two monkeys (118 IT neurons in M1 x 125 IT neurons in M2 = 14750 pairs in MUA). For each pair, we computed the firing rate of the two neurons in 10 ms bins, and calculated their cross-correlation within a time window of 2 s with a maximum lag of 1 s. The average cross-correlogram between all IT neuron pairs (n = 14,750) is shown in Figure S13A. It can be seen that the cross-correlation shows rich dynamics with time, although we did not observe any link to key behavioral events or grooming states.

To further characterize these cross-correlation dynamics, we first grouped the time bins into the three types of grooming behaviors (M1 grooming M2, M2 grooming M1 and no grooming), and calculated the cross-correlogram for each grooming type. The resulting cross-correlogram showed no clear differences across grooming type (Figure S13B). Next, we identified the lag at which there was a peak in the average cross-correlation. The resulting peak average cross-correlation (Figure S13C) varied with time, showing the largest cross-correlation for M2 grooming M1, followed by No Grooming, and then by M1 grooming M2 (Figure S13D). Finally, we also calculated the peak lag across time (Figure S13E) which did not show any systematic variation with grooming types (Figure S13F). Interestingly, both give and receiving grooming conditions had a positive lag, meaning that M2 activity precedes M1 activity in both conditions, for reasons that are not clear to us.

We repeated these analysis for PMv-PFC pairs (118 PMv/PFC neurons in M1 x 128 PMv neurons of M2 = 15104 pairs). We grouped PMv & PFC together so that there would be roughly equal numbers of neurons in M1 & M2, after finding the results to be qualitatively similar even without grouping. We observed less temporal variations in the average cross-correlogram compared to IT (Figure S13G), with no systematic differences between grooming bouts (Figure S13H) and in peak average correlations (Figure S13I-J). However with peak lags, we observed a negative mean lag in the M2 grooming M1 condition, indicating that M2 activity was lagging behind M1 (Figure S13L), consistent with our main finding that the receiver of grooming drives the social interaction.

Finally, we repeated these analyses using the LFP signal isolated from each electrode (106 M1 IT neurons x 109 M2 IT neurons = 11554 neuron pairs; 103 M1 PMvPFC neurons x 110 M2 PMv neurons = 11330 pairs; Figure S14). We subsampled the data (in 10 ms bins as done for MUA), for the ease of processing. Here too, we did not observe any systematic differences in cross-correlation dynamics between the different grooming types.

In sum, we conclude that there are rich dynamics of cross-correlation between both brains during social interaction, but these dynamics are not clearly interpretable at least to the best of our efforts.

**Figure S13.**
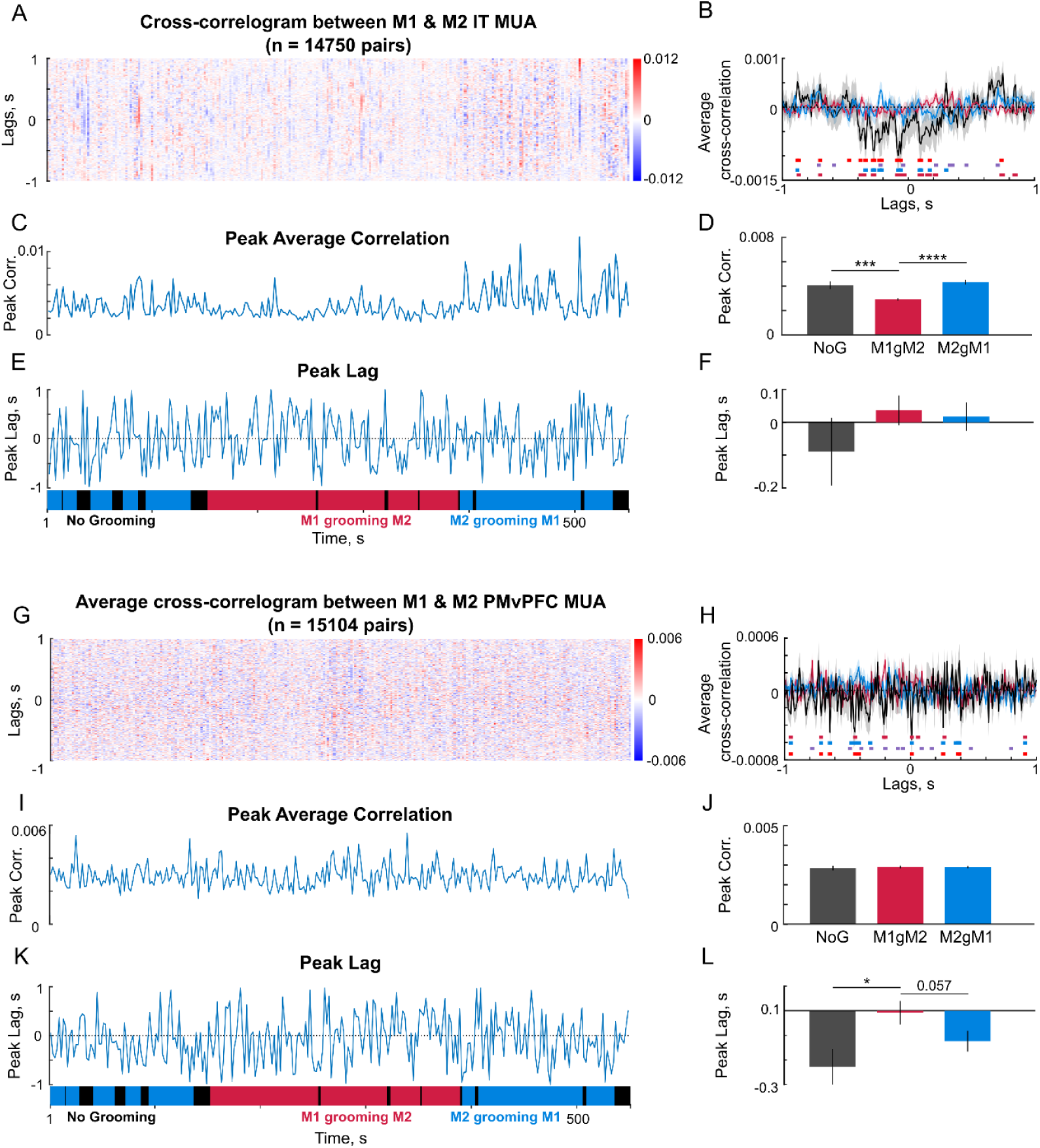
MUA cross-correlation dynamics during social interaction. (A) Colormap of average cross-correlogram at each lag across IT neuron pairs across time. Positive lags indicate M2 activity precedes M1 activity and negative lags indicate the opposite. (B) Average cross-correlogram in the three different grooming conditions. The horizontal lines indicate the lags at which the median cross-correlation differs significantly between various grooming conditions (p<0.05, rank sum test): Pink – M1gM2 vs No grooming, Blue – M2gM1 vs No grooming, Purple – M1gM2 vs M2gM1, Red – [M1gM2, M2gM1 combined] vs No grooming. (C) Peak of the average cross-correlation across time. (D) Average of the peak cross-correlation for each grooming type. Error bar indicate s.e.m. of the peak correlation across time bins. (E) Peak lag (time at which average cross-correlation attains a peak) across time. (F) Average peak lag for each grooming type. Error bars indicate s.e.m across time bins. (G-L) Same as (A-F) but for PMv/PFC neuron pairs.

**Figure S14.**
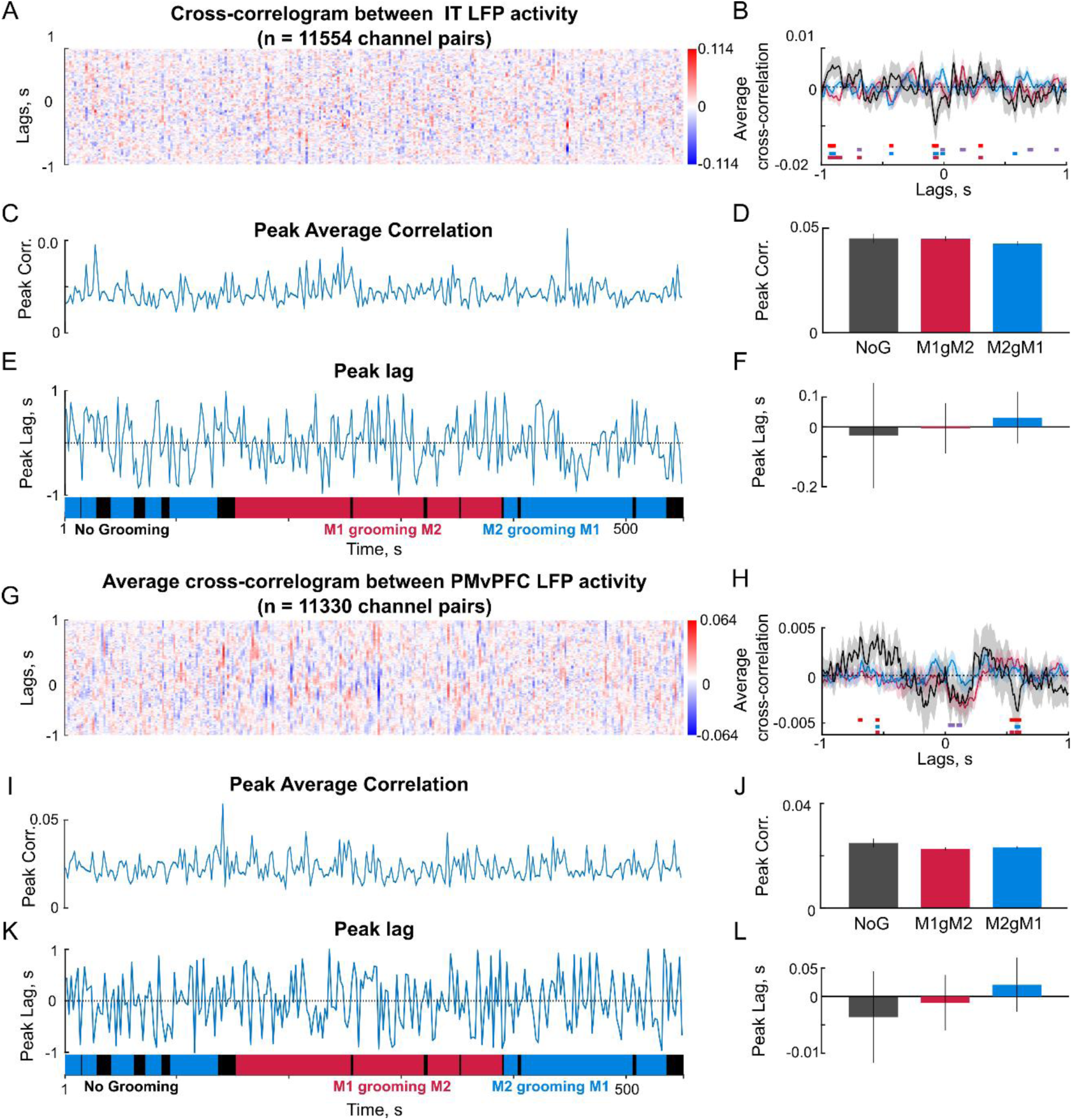
LFP cross-correlation dynamics during social interaction. (A) Colormap of average cross-correlation at each lag across IT electrode pairs across time. Positive lags indicate M2 activity precedes M1 activity and negative lags indicate the opposite. (B) Average cross-correlogram in the three different grooming conditions. The horizontal lines indicate the lags at which the median cross-correlation differs significantly between various grooming conditions (p<0.05, rank sum test): Pink – M1gM2 vs No grooming, Blue – M2gM1 vs No grooming, Purple – M1gM2 vs M2gM1, Red – [M1gM2, M2gM1 combined] vs No grooming. (C) Peak of the average cross-correlation across time. (D) Average of the peak cross-correlation for each grooming type. Error bar indicate s.e.m. of the peak correlation across time bins. (E) Peak lag (time at which average cross-correlation attains a peak) across time. (F) Average peak lag for each grooming type. Error bars indicate s.e.m across time bins. (G-L) Same as (A-F) but for PMv/PFC neuron pairs

#### Inter-brain coherence dynamics during social interaction

We next investigated whether there is interbrain coupling in specific frequency bands during the social interactions. Specifically we asked if there is a consistent phase relationship between the frequency content of neural firing rate or local field potentials, and whether there are any differences between the different grooming types.

#### Methods

We calculated the coherence between the MUA signal on every pair of electrodes in a given brain region, and averaged it across electrodes to get a global measure. We used MATLAB function “mscohere” to compute coherence, with the following set of parameters: Hamming Window of size 2048 samples, which comes to 0.82 s of data with a sampling rate of 2500 samples/s, and an overlap of 512 samples (0.25% overlap). This set of parameters gave a frequency resolution of 1.22 Hz. For the LFP signal, we used the same method as above to compute the coherence between every pair of neurons in a given brain region.

#### Results

We first visualized the coherence in the MUA signal between IT neuron pairs between the two monkeys across time in each brain region (Figure S15A). To assess whether the coherence is different for different grooming types, we averaged this coherence across time bins of each grooming type. This revealed significant differences in coherence at low frequencies between the different grooming conditions (Figure S15B). Since we did not observe differences in specific frequencies, we calculated the average coherence signal as a function of time after averaging across frequencies (Figure S15C). This revealed that there was significantly larger coherence during M1 grooming M2, followed by M2 grooming M1 with least coherence overall in the no-grooming condition (Figure S15D). We found similar results for PMvPFC neuron pairs as well (Figure S15E-H).

We performed a similar analysis on the LFP signal (Figure S16). Although there were differences in coherence at specific frequencies (Figure S16B,F), we did not find a systematic difference in coherence overall (Figure S16D,F).

We conclude that there is increased coherence at all frequency in multiunit activity during social interaction periods in all recorded regions. However, the variations in coherence with time are not clearly interpretable at least to the best of our efforts.

**Figure S15:**
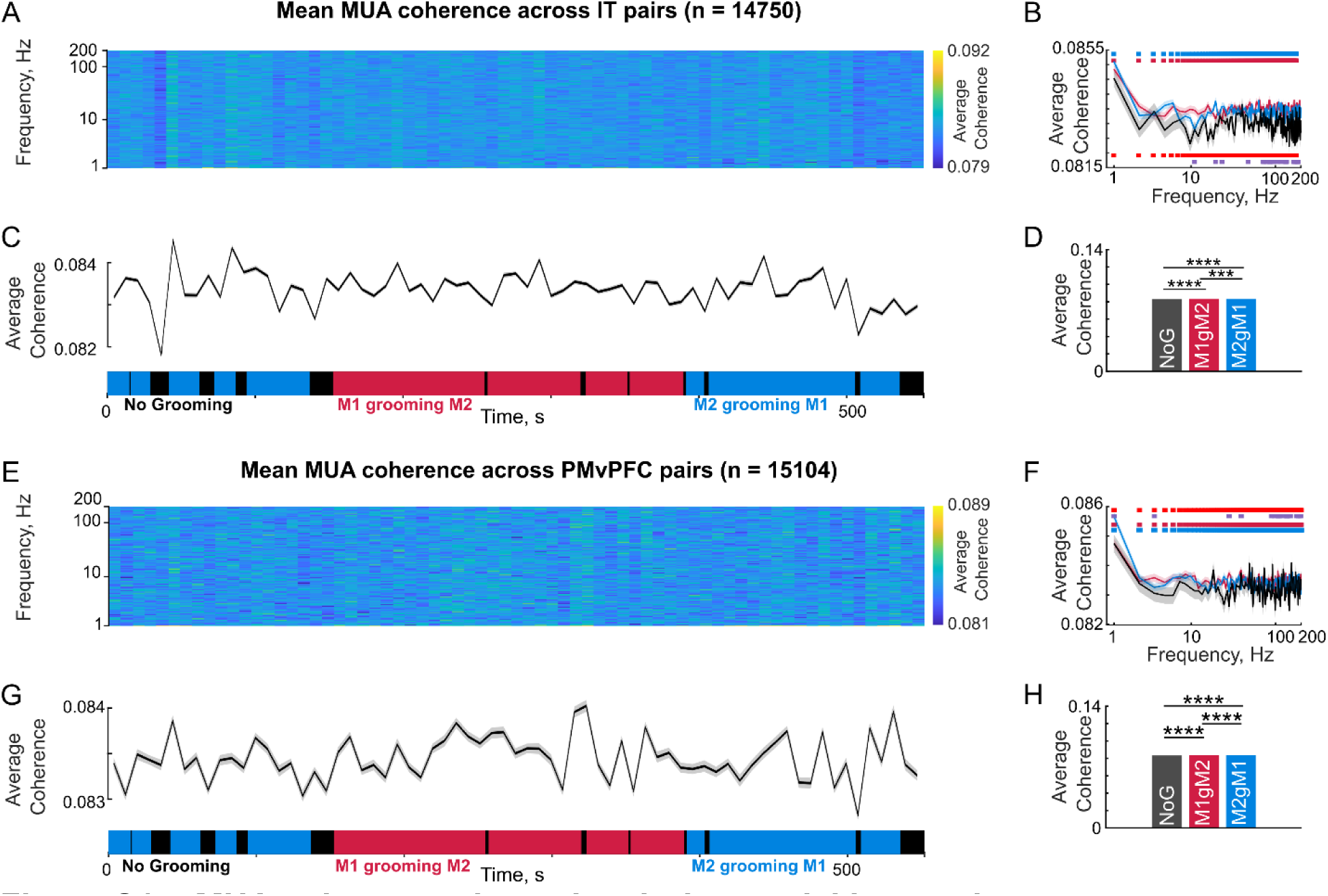
MUA coherence dynamics during social interaction. (A) Colormap of average magnitude-squared (ms) coherence across IT neuron pairs between the two monkeys (n = 14,750) for each frequency across time. (B) Average coherence for each frequency across grooming conditions (M1 grooming M2, M2 grooming M1 and No grooming). The horizontal lines indicate the frequencies at which the median coherence is significantly different between various grooming conditions(p<0.05, sign rank test): Pink – M1gM2 vs No grooming, Blue – M2gM1 vs No grooming, Purple – M1gM2 vs M2gM1, Red – [M1gM2, M2gM1 combined] vs No grooming. (C) Average coherence in low frequencies (0-200 Hz) across time. Shaded error bars indicate s.e.m across frequency. (D) Average coherence in low frequencies (0-200 Hz), for each grooming condition (M1 grooming M2, M2 grooming M1 and No grooming) across channel pairs. Asterisks indicate statistical significance of the comparisons (**** is p <0.00005, sign rank test across neurons). (E-H) Same as panels (A-D) but for PMvPFC pairs (n = 15,104).

**Figure S16.**
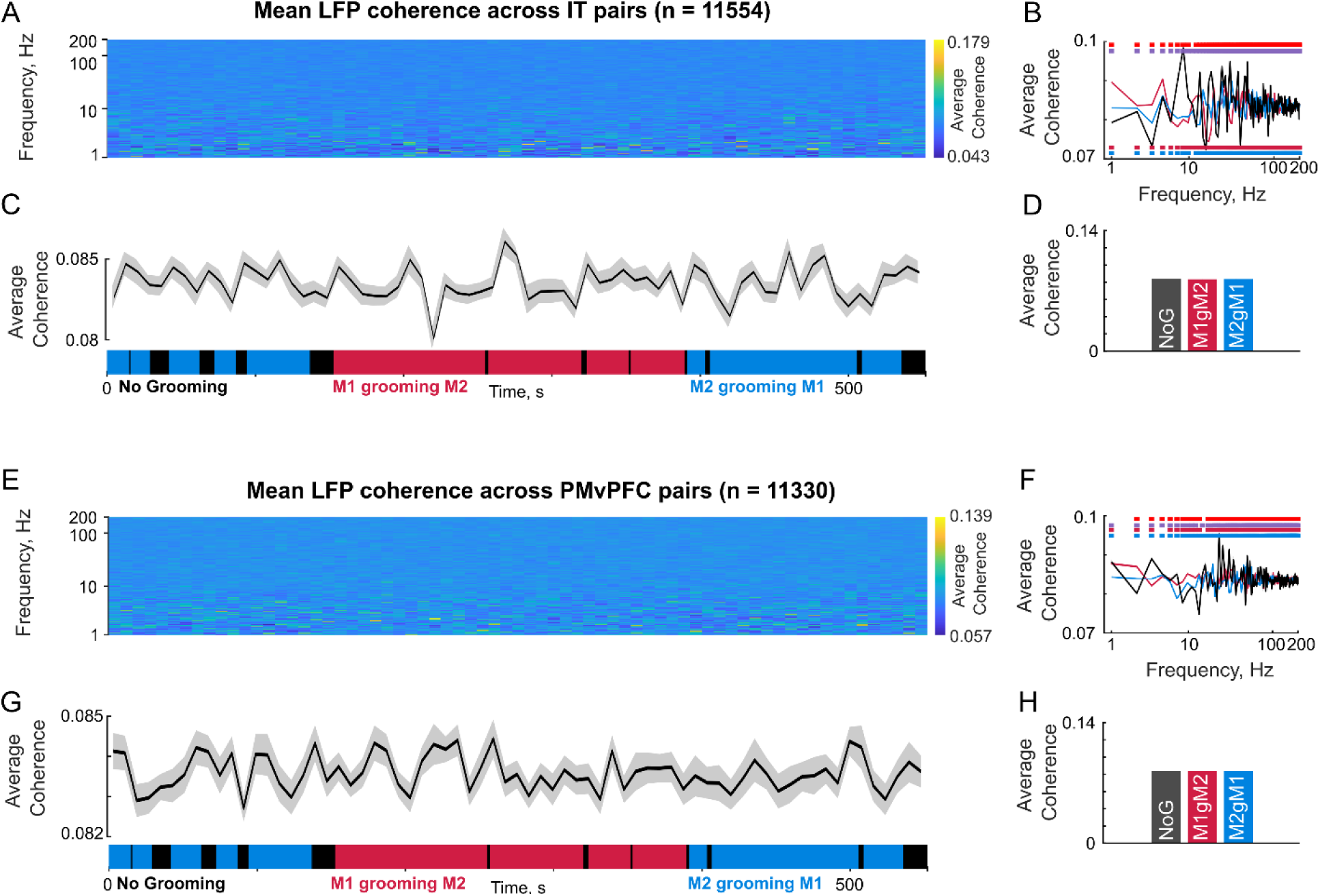
LFP coherence dynamics during social interaction. (A) Colormap of average LFP coherence of IT electrode pairs between the two monkeys (n = 11554) for each frequency across time. (B) Average coherence for each frequency across grooming conditions (M1 grooming M2, M2 grooming M1 and No grooming). The horizontal lines indicate the frequencies at which the median coherence is significantly different between various grooming conditions(p<0.05, sign rank test): Pink – M1gM2 vs No grooming, Blue – M2gM1 vs No grooming, Purple – M1gM2 vs M2gM1, Red – [M1gM2, M2gM1 combined] vs No grooming. (C) Average coherence in low frequencies (0-200 Hz) across time. Shaded error bars indicate s.e.m across frequency. (D) Average coherence in low frequencies (0-200 Hz), for each grooming condition (M1 grooming M2, M2 grooming M1 and No grooming) across channel pairs. (E-H) Same as panels (A-D) but for PMvPFC pairs (n = 11330).

### SECTION S9. CONVERGENCE BETWEEN BEHAVIOR AND NEURAL SIGNALS

#### Summary

In the main text, we have shown that neural signals like grooming surplus, partner identity, partner joints, interbrain coupling vary dynamically across the social interaction, and are modulated by the grooming state. However it is possible that key behavioral events during the social interaction have converging evidence from multiple neural signals, which might support a clear indication of each monkeys’ internal state.

To this end, we carefully reviewed the behavioral events throughout the social interaction, and identified key events that we considered potentially important. At those times, we report the status of several neural signals that we considered to be relevant to the internal state of each monkey. These neural signals included

1. Alpha power, known to be a marker for relaxation or eye closed
2. Beta power, generally thought to be a marker for movements
3. Partner decoding accuracy from M1 & M2 IT neurons (same as Figure S7), which could indicate the degree to which each monkey is aware of his social partner
4. Joint model correlations for self & partner joints for M1 & M2, which could indicate the degree to which each monkey is tracking his own and his partner’s joints
5. Partner model correlation for M1 & M2, which could indicate the degree to which the two brains are coupled.

**Table S1.** Key behavioral events and ongoing neural signals (ref: Figure S17 & Video S2)

| Video Time | Key Behavioral Event | Summary of Ongoing Neural Signals |
| --- | --- | --- |
| 145 s | M2 was grooming M1 and changed the grooming site without no obvious indication from M1. | • *Alpha power*: No clear trend.<br>• *Beta power*: Peak in M1 IT.<br>• *Partner decoding*: High in M1 and M2.<br>• *Joint models*: Self & partner model high in M1, self model high in M2.<br>• *Partner neural model*: M2 > M1<br>• *Interpretation*: Both M1 & M2 are tracking their partner's identity & joints. |
| 159 to 162 s | M2 stopped grooming M1 with no obvious indication from M1<br>M1 was lying down and sat up as soon as M2 stopped, but did not look directly at M2. | • *Alpha power*: No clear trend.<br>• *Beta power*: Peak in M1 PMv & M2 IT, PMv.<br>• *Partner decoding*: High in M1, Low in M2<br>• *Joint models*: M1 & M2 self models are high.<br>• *Partner neural model*: M2 > M1.<br>• *Interpretation*: M1 is aware of M2 despite not looking directly, and M1 & M2 are not tracking their partner's joints. |
| 168 to 172 s | M2 is not grooming M1<br>M1 made a sound (presumably soliciting grooming)<br>M2 responded by resuming grooming of M1. | • *Alpha power*: Peak in M1 PMv.<br>• *Beta power*: Peak in M1 PMv & M2 IT.<br>• *Partner decoding*: M1 > M2.<br>• *Joint models*: self > partner in M1, partner > self in M2, and both trends reduced over time. |
|  |  | • *Partner neural model*: M2 > M1, but both are at a peak.<br>• *Interpretation*: M1 is aware of his partner's identity, and both M1 & M2 are tracking their partner's joints. |
| 192 to 202 s | M2 was grooming the right side of M1's face, then switched hands to hold M1's implant and bend M1's head to groom the left side of M1's face. | • *Alpha power*: No clear trend.<br>• *Beta power*: No clear trend.<br>• *Partner decoding*: M2 > M1.<br>• *Joint models*: No clear trend.<br>• *Partner neural model*: Small peak in M1 & M2.<br>• *Interpretation*: M2 is directly seeing his partner and encoding his partner's identity, and both M1 & M2 are encoding their own joints and their partner's joints. |
| 216 to 220 s | M2 was grooming M1's back and stopped grooming M1 momentarily.<br>M1 turned, looked up, stretched his neck and made a small movement (presumably soliciting grooming)<br>M2 responded by resuming grooming of M1 starting with the neck. | • *Alpha power*: Peak in M1 IT, PMv, PFC.<br>• *Beta power*: Peak in M1 IT, PMv, PFC & M2 IT.<br>• *Partner decoding*: M1 > M2<br>• *Joint models*: self & partner high in M1, partner > self in M2.<br>• *Partner neural model*: M1 > M2<br>• *Interpretation*: M1 is aware of his partner's identity and his own & partner's joints. M2 is tracking his partner's joints. |
| 266 to 282 s | M2 stopped grooming M1, and M1 turned around and began scratching his leg<br>M2 turned around, looked down at M1 and then presented his back to M1 (presumably soliciting grooming)<br>M1 huddled against M2 and both made lip-smacking gestures<br>M1 then started grooming M2. | • *Alpha power*: Peaks in M1 IT, PFC and M2 IT.<br>• *Beta power*: M1 beta power declining in IT, PMv, PFC and peaks in M2 IT.<br>• *Partner decoding*: M1 > M2 at first, then M2 > M1<br>• *Joint models*: Partner > self for M1, no clear trend for M2.<br>• *Partner neural model*: M1 > M2<br>• *Interpretation*: Initially M1 tracked his partner's identity and then M2 tracked his partner's identity. M1 is closely tracking his partner's joints, while M2 is tracking both his own & partner's movements. M1's brain activity is better predicted by M2, again indicating coupling. |
| 385 to 388 s | M1 was grooming M2's face. M2 turned around to present his left shoulder to M1, and M1 responds by grooming there. | • *Alpha power*: Peak in M1 IT & PFC.<br>• *Beta power*: Peak in M1 IT & PFC.<br>• *Partner decoding*: Peak in M2<br>• *Joint models*: No clear trend but partner > self soon afterward in M2, and self > partner in M1.<br>• *Partner neural model*: No clear trend but M1 > M2 soon afterwards.<br>• *Interpretation*: M1 is relaxed (based on alpha power), M2 is aware of his partner's identity, and both monkeys are tracking the groomer's |
|  |  | joint movements. M1 brain activity is predicted by M2, indicating coupling. |
| 449 to 451 s | M1 was grooming M2's back<br>M2 turned around to present his chest to M1, and M1 responds by grooming there. | • *Alpha power*: Peak in M2 IT.<br>• *Beta power*: Peak in M2 IT.<br>• *Partner decoding*: M2 > M1<br>• *Joint models*: No clear trends.<br>• *Partner neural model*: M1 > M2<br>• *Interpretation*: M2 is tracking his partner's identity, and M1's brain is predicted by M2, indicating coupling. |
| 480 to 482 s | M1 was grooming M2's right shoulder and stopped and looked downwards and touched M2's leg.<br>M2 extended his right leg, presumably in response.<br>M1 started grooming M2's right leg. | • *Alpha power*: Peak in M2 IT & PMv<br>• *Beta power*: Peak in M2 IT & PMv<br>• *Partner decoding*: M1 > M2<br>• *Joint models*: self > partner in both monkeys.<br>• *Partner neural model*: M1 > M2<br>• *Interpretation*: M1 is tracking his partner's identity, M1 & M2 are tracking their own joints, but M1's brain is predicted by M2, indicating coupling. |
| 520 s | M1 was grooming M2's foot.<br>M2 starts grooming M1.<br>M1 responds by stopping his grooming of M2. | • *Alpha power*: Peak in M1 IT, PMv, PFC soon afterwards.<br>• *Beta power*: Peak in M1 IT, PMv, PFC soon afterwards.<br>• *Partner decoding*: M2 > M1<br>• *Joint models*: partner > self for M2 and self > partner for M1.<br>• *Partner neural model*: M2 > M1 soon afterwards.<br>• *Interpretation*: M1 seems to relax (based on alpha power), M2 is aware of his partner's identity, and both monkeys are tracking the receiver's joints. M2's brain is predicted by M1, indicating coupling. |
| 533 s | M2 is grooming M1 and making small changes in the groomed locations on M1's back without any obvious indications from M1. | • *Alpha power*: Peak in M1 IT, PMv, PFC.<br>• *Beta power*: Peak in M1 IT, PMv, PFC.<br>• *Partner decoding*: M2 > M1<br>• *Joint Models*: partner > self in M1, self > partner in M2.<br>• *Partner neural model*: M2 > M1<br>• *Interpretation*: M1 is relaxed (based on alpha power), M2 is tracking his partner's identity, both monkeys are tracking the groomer's joints, M2's brain is predicted by M1, indicating coupling with receiver's brain. |
| 570 to 572 s | M2 was grooming M1's middle back.<br>M1 then lay down on the ground<br>M2 responds by grooming M1's lower back. | • *Alpha power*: Peak in M1 IT, PMv, PFC<br>• *Beta power*: Peak in M1 IT, PMv, PFC<br>• *Partner decoding*: M2 > M1<br>• *Joint models*: partner > self in M2 at first, then partner > self in M1. |
|  |  | • *Partner neural model*: M1 > M2<br>• *Interpretation*: M1 is relaxed, M2 is tracking his partner's identity, M2 is tracking his partner's joints and then M1 is tracking his partner's joints, M1's brain is predicted by M2, indicating coupling with the groomer/dominant partner's brain. |
| 625 to 628 s | M2 was grooming M1's lower back<br>M1 starts touching his implant and licking his hand.<br>M2 looks up to see what M1 is doing. | • *Alpha power*: No clear trend.<br>• *Beta power*: No clear trend.<br>• *Partner decoding*: low in M1 & M2.<br>• *Joint models*: self > partner in M1, partner > self in M2.<br>• *Partner neural model*: M1 > M2<br>• *Interpretation*: M2 is tracking M1's joints to see what M1 is doing. |
| 633 to 638 s | M2 stopped grooming M1, who was lying down.<br>M1 responds by getting up and turning to present his neck<br>M2 responds by grooming M1's neck. | • *Alpha power*: Peak in M1 IT, PMv, PFC and in M2 IT.<br>• *Beta power*: Peak in M1 PMv soon afterwards, Peak in M2 IT & PMv.<br>• *Partner decoding*: M2 > M1<br>• *Joint models*: self > partner at first in M1, then partner > self, partner > self in M2.<br>• *Partner neural model*: M2 > M1<br>• *Interpretation*: M1 is relaxed and M2 is relaxed, M2 is tracking his partner's identity, M1 is moving but also tracking his partner's joints, M2 is tracking his partner's joints, M2's brain is predicted by M1, indicating coupling with receiver's brain. |
| 640 to 645 s | M2 holds M1's implant and bends M1's head to groom his right temple. | • *Alpha power*: Peak in M1 IT & PFC<br>• *Beta power*: Peak in M1 PMv & later in M1 IT & PFC.<br>• *Partner decoding*: M2 > M1<br>• *Joint model*: No data, since a few joints were not visible in the top view.<br>• *Partner neural model*: M2 > M1<br>• *Interpretation*: M1 is relaxed, M2 is tracking his partner's identity, M2's brain is predicted by M1, indicating coupling with receiver's brain. |

**Table S2.** Key neural events and the accompanying behaviors (ref: Figure S17 & Video S2)

| Video Time | Summary of Ongoing Behavior | Key Neural Event |
| --- | --- | --- |
| 179 to 184 s | M2 was grooming M1's head<br>M1 closed his eyes. | Key Neural Event<br>• *Alpha power*: Peak in M1 IT, PMv, PFC.<br>• *Beta power*: Peak in M1 IT, PMv, PFC.<br>Other Neural Signals<br>• *Partner decoding*: Drops for M1<br>Interpretation<br>• *M1 is relaxed. Closing eyes is well-known to cause increased alpha power.* |
| 235 to 260 s | M1 was looking up for a long time while M2 was grooming his neck. | Key Neural Event<br>• *Alpha power*: Peak in M1 IT, PMv, PFC.<br>• *Beta power*: Peak in M1 IT, PMv, PFC.<br>Other Neural Signals<br>• No clear trends.<br>Interpretation<br>• M1 is relaxed. |
| 346 to 360 s | M2 looked at the roof while M1 was grooming his chin/neck. | Key Neural Event<br>• *Alpha power*: Peak in M2 IT<br>• *Beta power*: Peaks in M1 IT, PFC & M2 IT<br>Other Neural Signals<br>• *Partner decoding*: M2 > M1<br>Interpretation<br>• M2 is relaxed but aware of his partner. |

**Figure S17.**
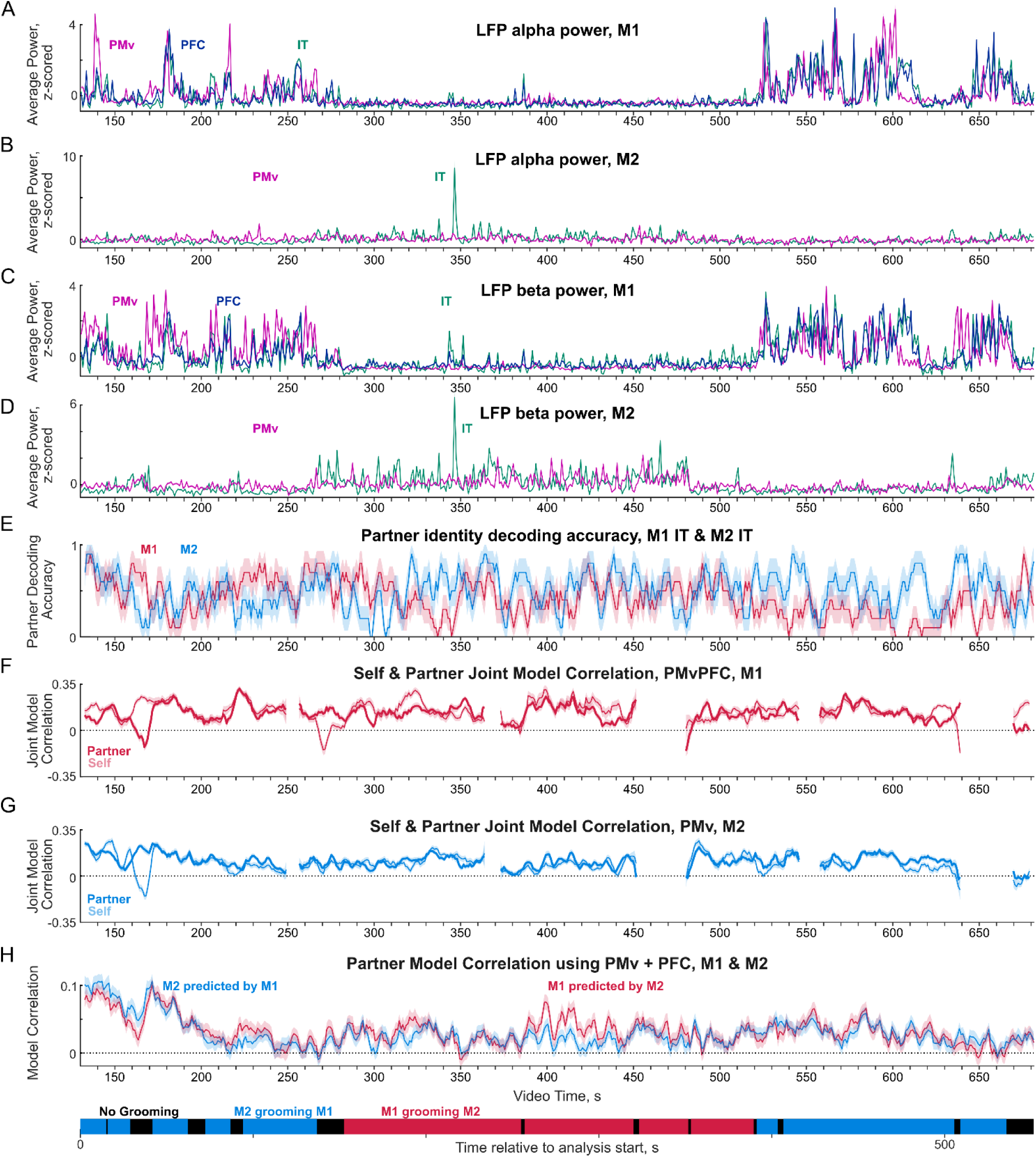
Convergent modulation of key behavioral & neural events (see Video S2) (A) Time course of average LFP power in the alpha band (8-12 Hz) for IT sites *(green)*, PMv sites *(purple)* and PFC sites *(blue)* in M1. (B) Same as (A) for M2. (C) Same as (A) but for beta band (12-30 Hz) in M1 sites. (D) Same as (C) for M2 sites. (E) Partner identity decoding accuracy, calculated in 5 s bins with a step size of 1 s from M1 & M2 IT (same as Figure S7). (F) Instantaneous correlation (smoothed with a 5 s moving window with a step of 1 s) of the partner (*dark*) and self (*light*) joint models across PMv & PFC sites for M1. All conventions are as in Figure S11. (G) Same as (F) for M2 PMv. (H) Time course of smoothened partner model correlations for M1 & M2 during the entire interaction (same as Figure 6D).

## SUPPLEMENTARY VIDEOS

**[Video S1.](https://osf.io/fdjwa) Video snippets demonstrating how the grooming receiver drove the social interaction.**

**[Video S2.](https://osf.io/nfpr5) Complete video of the social interaction session**

## Notes

https://osf.io/tu5ed

### Competing Interest Statement

The authors have declared no competing interest.

### Summary of Updates

Added a new supplementary figure and updated an existing supplementary figure(FigureS02)

